# Genome-wide quantitative trait loci mapping on *Verticillium* wilt resistance in 300 chromosome segment substitution lines from *Gossypium hirsutum* × *Gossypium barbadense*

**DOI:** 10.1101/788901

**Authors:** Md Harun or Rashid, Peng-tao Li, Tingting Chen, Koffi Kibalou Palanga, Wan-kui Gong, Qun Ge, Ju-wu Gong, Ai-ying Liu, Quan-wei Lu, Latyr Diouf, Zareen Sarfraz, Muhammad Jamshed, Yu-zhen Shi, You-lu Yuan

**Affiliations:** State Key Laboratory of Cotton Biology, Key Laboratory of Biological and Genetic Breeding of Cotton, The Ministry of Agriculture, Institute of Cotton Research, Chinese Academy of Agricultural Sciences, Anyang 455000, Henan, China; Senior Scientific Officer, Breeding Division, Bangladesh Jute Research Institute, Dhaka-1207, Bangladesh; School of Biotechnology and Food Engineering, Anyang Institute of Technology, Anyang 455000, Henan, China

**Keywords:** CSSLs, *Verticillium* wilt, Disease Index, Quantitative Trait Loci, Meta-analysis

## Abstract

Cotton *Verticillium* wilt (VW) is a devastating disease seriously affecting fiber yield and quality, and the most effective and economical prevention measure at present is selection and extension of *Gossypium* varieties harboring high resistant VW. However, multiple attempts to improve the VW resistance of the most widely cultivated Upland cotton have brought in little significant progress, and it seems necessary and urgent to develop Chromosome segment substitution lines (CSSLs) for merging the superior genes related with high yield and wide adaptation from *G. hirsutum* and VW resistance and excellent fiber quality from *G. barbadense*. In this study, 300 CSSLs were chosen from the developed BC_5_F_3:5_ CSSLs constructed by *G. hirsutum* CCRI36 and *G. barbadense* Hai1 to conduct quantitative trait locus (QTL) mapping on VW resistance, and a total of 53 QTLs relevant to VW disease index (DI) were identified together with the phenotypic data of 2 years investigations in two fields with two replications per year. All the QTLs were distributed on 20 chromosomes with phenotypic variation of 3.74-11.89%, of which 29 stable ones were consistent in at least two environments. Based on Meta-analysis on the 53 QTLs, 43 novel ones were identified, while 10 ones consistent to previously identified QTLs. Meanwhile, 32 QTL hotspot regions were detected, including 15 ones were novel. This study concentrates on QTL identification and screening hotspot region related with VW in the 300 CSSLs, which lay a solid platform not only for revealing the genetic and molecular mechanisms of VW resistance, but also for further fine mapping, gene cloning and molecular designing in breeding program for resistant cotton varieties.

## 1. Introduction

Cotton (*Gossypium* spp. L.) is not only the most significant cash crop producing the main source of natural fiber for the textile industry, but also the second important oilseed crop [1]. The cultivation history of cotton could retrospect to 7000 years ago[2], which is widely grown in approximately 100 countries principally located in tropical and sub-tropical arena [3]. The genus *Gossypium* consists of 53 species all over the world, including 46 diploid ones (2n = 2× = 26) and 7 allotetraploid ones (2n = 2× = 52) [4], of which the emergence of the latte dated from a polyploidization event between A and D genome1-2 million years ago [3]. Only 4 cultivated species (2 diploids and 2 tetraploids) are extant and widely planted all over the world, while the rest of the 53 species are wild but important reservoir of beneficial agronomic traits for improvement of the cultivated ones [5, 6]. Nowadays, *G. hirsutum* and *G. barbadense* are the the most widely cultivated species, and could contribute for 97% and 3% of world cotton production, respectively, which attributes to the facts that the former harbors high yield and wide adaptability, while the latter possesses superior fiber quality and high VW resistance [7].

Plenty of restraining factors during organism growth are generally divided into abiotic and biotic stresses [8], while plant diseases might be the dominating threat in cotton production [9], of which *Verticillium* wilt (VW) infected by soil-borne fungus *Verticillium dahliae* Kleb has been the most significant disease in cotton production due to causing substantial yield loss and serious fiber quality reduction [10–12]. As a result of cotton VW infestation, fiber loss is estimated to approximately stand at 80% [13]. What is worse, this disease can attack more than 400 plant species and exist in soil for a long period in dormant form in the vascular system of perennial plants. Thus, it is completely impossible to control VW disease through conventional method [14]. The general symptoms of the disease are vascular browning, stunting, leaf epinasty and chlorosis, curling or necrosis, wilt and finally death of the entire plant [15, 16].

Despite multiple methods put forward to control VW, it remains one of the most efficient and economical measures to develop elite cotton cultivars harboring genetic factors tolerant or completely resistant against pathogen in cotton breeding [17–19]. There are only four subsistent cultivars of *Gossypium* species, while the tetraploid cultivars cover more than 95% of planting areas around the world, namely as *G. barbadense* (Sea Island cotton) and *G. hirsutum* (Upland cotton), which present resistant and susceptible to VW disease, respectively [20, 21]. Hybrid breeding via conventional techniques has been utilized earlier to improve VW resistance in upland cottons, while some hindrances like infertility and hybrid break down/low parent heterosis hindered the way of conducting resistant gene introgression from *G. barbadense* into *G. hirsutum* [21]. Therefore, it has become a challenging task for cotton breeders to achieve synchronous improvement in cultivating novel varieties simultaneously harboring high yield, superior fiber quality, and high disease resistance. QTL Mapping approaches make it possible for the discovery of quantitative genetic factors responsible for disease resistance as well as high fiber quality and yield with the utilization of marker-assisted selection (MAS). Thus, we can take full advantage of genetic markers presenting linkage disequilibrium with disease resistance to confirm the contribution of key candidate genes in cotton research, which will be transferred from Sea Island cotton into Upland cotton to improve the VW resistance [22].

Chromosome Segment Substitution Lines (CSSLs) have perpetual effects as accompanied with similar genetic base to their recurrent parent thereby acting as favorable implement in mining of elite QTLs and alleles; ultimately carrying out advanced functional genomic techniques devoid of any non-additive genetic effects [23–28]. Optimal utilization of upland cotton as well as island cottons can be brought about via MAS and conventional breeding techniques of inbreeding, outcrossing and backcrossing with the provision of CSSLs. Therefore, CSSLs are extensively exploited especially in QTL mapping approaches for discovering genetic factors responsible for economic traits such as fiber quality, yield, biotic and abiotic stress tolerance or resistance [29–37].

Nowadays, cotton genomics research like other crop species, has been successively performed by QTL mapping on the significant traits based upon comprehensive deployment of molecular markers, of which simple sequence repeats (SSRs) are the most extensively utilized genetic markers in cotton [38]. To date, approximately 19010 SSRs have been accounted for cotton genomics research in Cotton Data Base (http://cottondb.org/), and almost 100,290 microsatellites have been newly extracted from genome while about 77,996 ones have been established successfully.

In the recent days, there is a newly emerging technique of mapping renowned as Meta-analysis of QTLs in tetraploid cotton research, which has been intensively activated for the identification of hotspot regions and known to harbor a massive amount of QTLs [32, 33]. Consensus map positions for QTLs and merging of datasets are the fundamental properties for meta-analysis approach, making this technique unique and widely adoptable. Not only previously declared QTLs positions can be reassured with identification of hotspot regions, but also the pleotropic effects of QTLs for different traits can be identified with Meta QTL analysis [32]. Moreover, this beneficial aspect of meta-analysis can be exploited to create hotspot region refuging stable QTLs for any disease by reassembling the previously identified QTLs for the relevant disease. Facilitation of breeders and geneticists can be brought about by employing this technique as they would only need to identify that specific chromosome region enriched with genetic factors controlling disease resistance for MAS or advanced mapping techniques [7, 39].

The goals of this study therefore are to identify favorable QTL alleles linked with VW resistance, to screen SSR markers that can be implemented in marker-assisted breeding program, and to confirm consistent and stable QTLs through meta-analysis for MAS application in cotton breeding for VW prevention and control. The results in this study are of importance for VW resistance as well as breeding improvements in cotton.

## 2. Results

### 2.1 Phenotypic disease index (DI) of parents and controls

At Anyang in July 2015, the highest DI value of VW was obtained in the susceptible Jimian11 (41.95%), followed by CCRI36 (31.03%), while the lowest one was observed in the parental line Hai1 (6.21%) (Table 2), indicating a significant difference of DI values between Hai1 and Jimian11. At Anyang in August 2015, the highest DI was found in Jimian11 (48.30%), followed by CCRI36 (47.70%) and by Hai1 (19.50%). The difference of DI values between the parental lines was significant while that of DI values between CCRI36 and Jimian11 was insignificant (Figure 1. A). In both case of Xinjiang in July and August 2015, highly significant differences were observed between parental lines (Figure 1. B).

**Figure 1.**
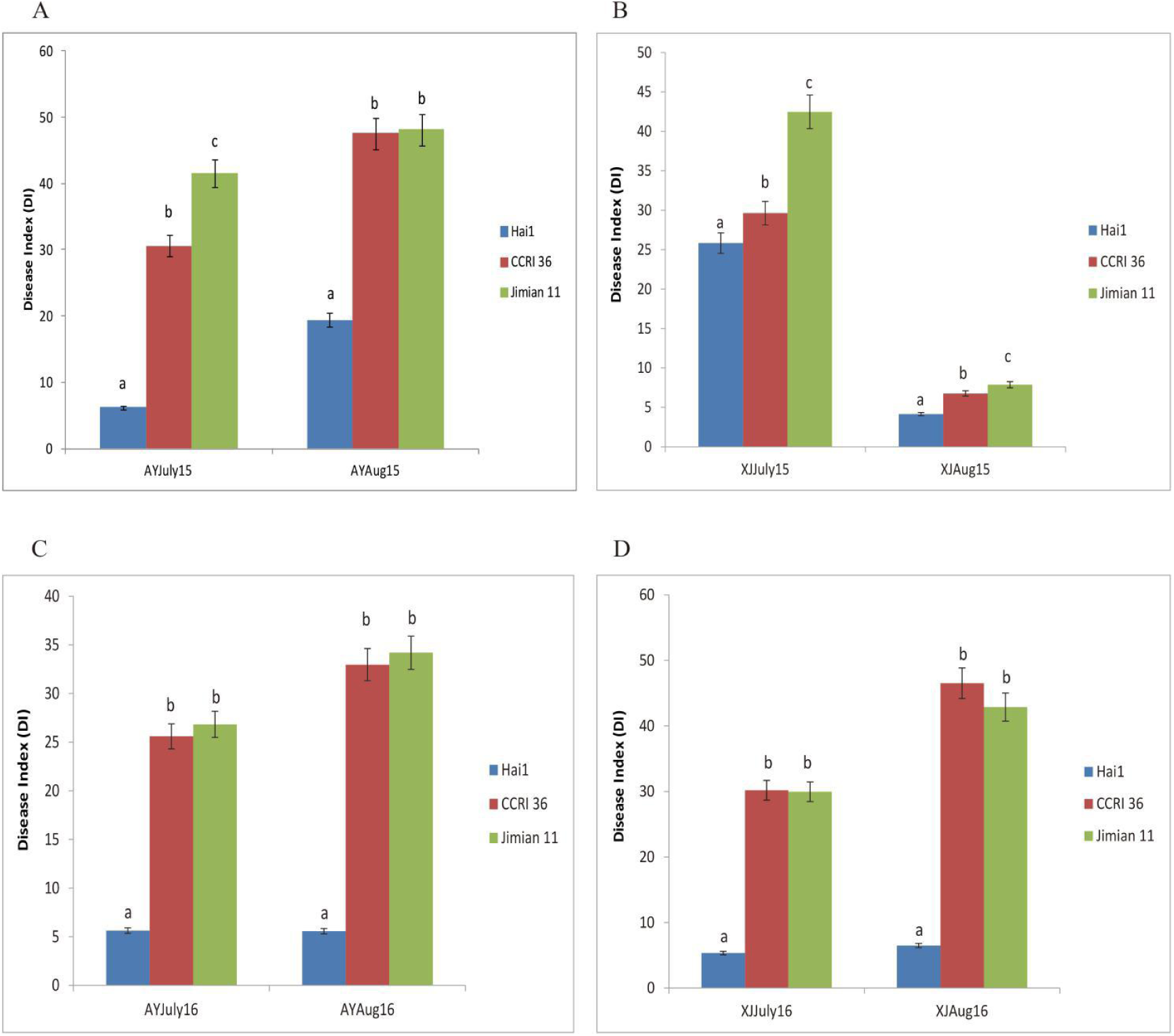
*Verticillium* wilt disease index of parent CCRI36, resistant control Hai1 and susceptible control Jimian11; Anyang 2015; (B) Xinjiang 2015; (C) Anyang 2016; (D) Xinjiang 2016. The error bar shows the standard deviation. a, b, c indicate the significance at 5%.

At Anyang in July 2016, the DI value of Jimian11 (26.83%) was the highest, followed by CCRI36 (25.57%), while the DI value of Hai1 (5.59%) was the lowest (Table 2), identifying no significant difference of DI values between CCRI36 and Jimian11. At Anyang in August 2016, the highest DI was recorded in Jimian11 (35.19%), followed by CCRI36 (32.89%), while the DI value of Hai1 (5.60%) was the lowest (Figure 1. C). The difference of DI values between CCRI36 and Jimian11 was also insignificant. In both case of Xinjiang in July and August 2016, we observed highly significant difference of resistance against the VW disease between the parents, while no significant difference between CCRI36 and Jimian11 was observed (Figure 1. D).

### 2.2 Evaluation of CSSLs for VW resistance

The ANOVA results displayed the P-value was 0.002, suggesting significant differences of resistance against VW in CSSLs (Table 1). Results of the descriptive statistical analysis of CSSLs and parental lines across 8 environments were illustrated in Table 2. Less than one absolute value of skewness of the mean values of VW in CSSLs across 8 environments indicated a normal distribution. The DI of CSSLs presented a perpetual and normal distribution, which was in consistent with multi-gene inheritance patterns for VW resistance (Figure 2).

**Figure 2.**
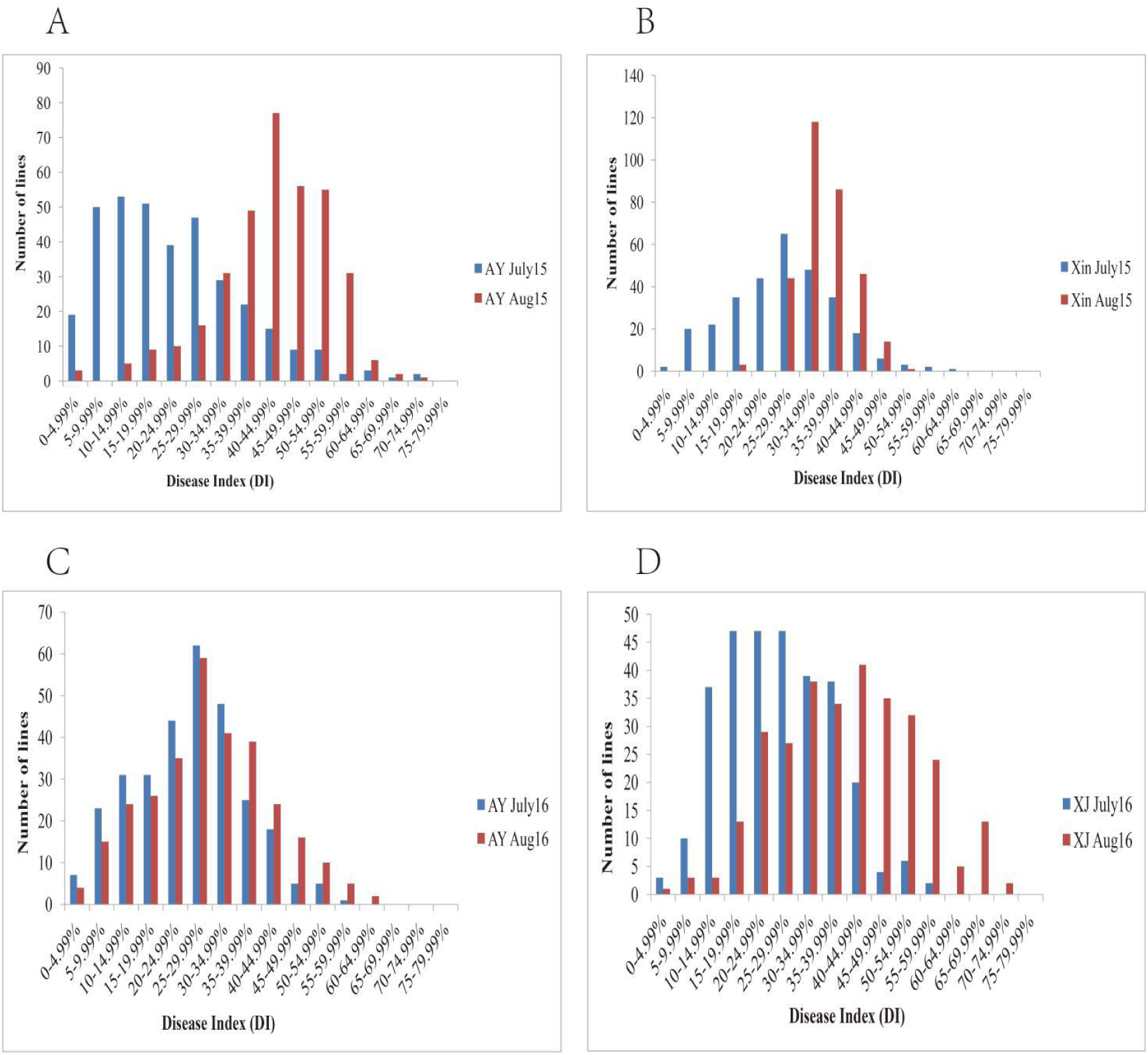
Normal distribution of DI phenotype in CSSLs; (A) Anyang 2015; (B) Xinjiang 2015; (C) Anyang 2016; D Xinjiang 2016.

**Table 1.** Analysis of variance of VW resistance ratings showed by DI across 8 environments

| Source of Variation | DF | Sum of square | Mean square | F | P-value |
| --- | --- | --- | --- | --- | --- |
| Environments | 7 | 281489 | 40212.71 | 508.556 | 0** |
| Genotypes | 299 | 101795.6 | 340.4534 | 4.306 | <0.001** |
| Error | 2093 | 165498.4 | 79.07235 |  |  |
| Total | 2399 | 548783 |  |  |  |

**Table 2.**
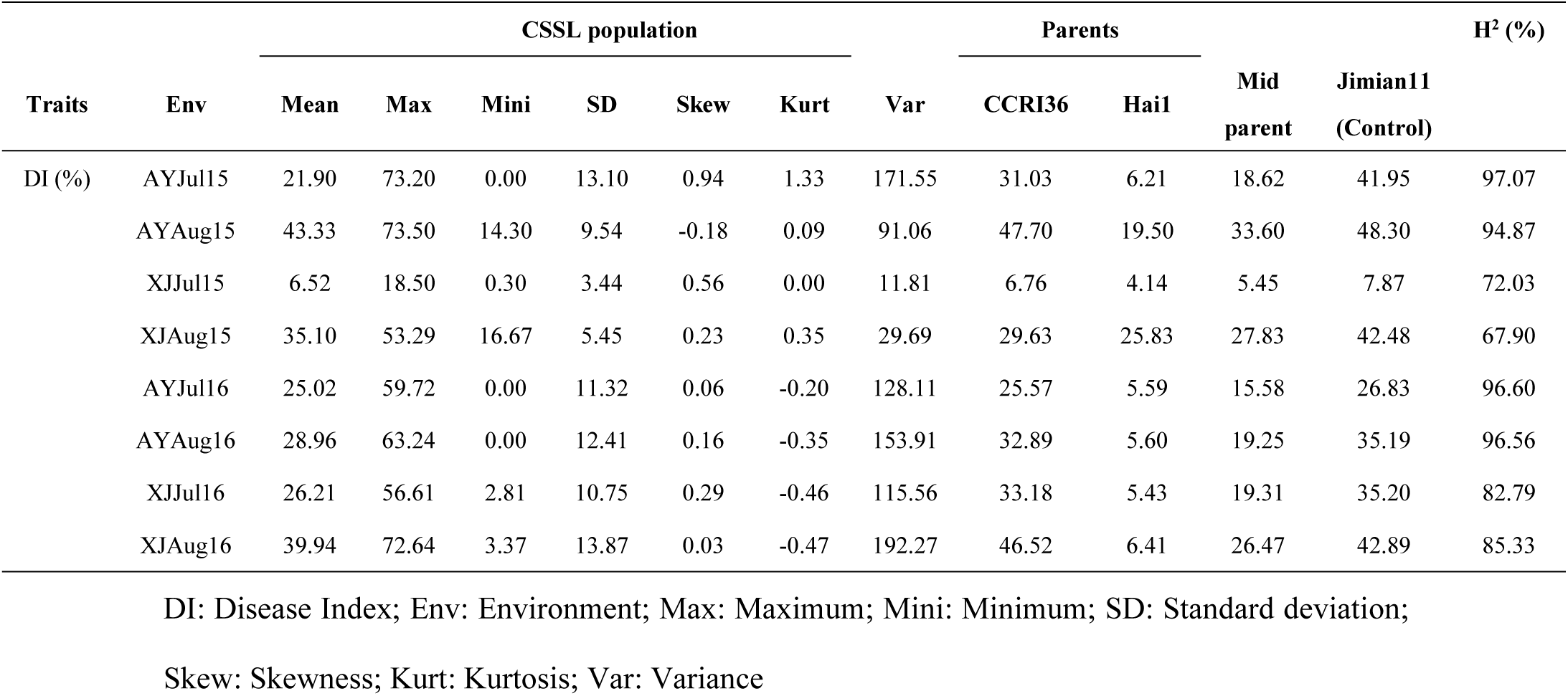
Descriptive statistics of VW resistance with broad sense Heritability (H^2^) measured in

The average DI values of CSSLs varied from 0.30 to 18.50% in XJJuly15 and from 16.67 to 53.29% in XJAug15 (Table 2). The average DI value in XJJuly15 was 6.52%, showing not significant to either of parents. On the other hand, the average DI values of CSSLs varied from 0 to 59.72% in AYJuly16. The average DI value in AYJuly16 was 25.02%, which was close to the recurrent parent CCRI36 (25.57%). The broad-sense heritability varied from 67.90% to 97.07%, of which the highest heritability was observed in AYJuly15 while the lowest in XJAug15 (Table 2). For all the environments of two years and developmental stages, wide variations of heritability were found in CSSLs to VW disease onset with some lines showing introgressive segregation over their parents.

### 2.3 Correlation coefficient among DI in different stages growth and environments

Highly significant positive correlations were visible among the disease index of *Verticillium* wilt in the fields except between XJJul15 and AYJul16 (Table 3).

**Table 3.** Correlation coefficient among the DI in the different stages of growth of BC5F3:5 population

| Traits | AYJul15 | AYAUG15 | XJJul15 | XJAug15 | AYJul16 | AYAUG16 | XJJul16 |
| --- | --- | --- | --- | --- | --- | --- | --- |
| AYAUG15 | 0.407** |  |  |  |  |  |  |
| XJJul15 | 0.202** | 0.164** |  |  |  |  |  |
| XJAug15 | 0.187** | 0.136* | 0.314** |  |  |  |  |
| AYJul16 | 0.119* | 0.164** | 0.04 | 0.123* |  |  |  |
| AYAUG16 | 0.315** | 0.326** | 0.169** | 0.188** | 0.401** |  |  |
| XJJul16 | 0.445** | 0.485** | 0.210** | 0.157** | 0.248** | 0.376** |  |
| XJAug16 | 0.437** | 0.481** | 0.163** | 0.164** | 0.240** | 0.379** | 0.919** |

### 2.4 QTL mapping

In total, 53 QTLs for VW were detected during different stages of growth and environments at Anyang and Xinjiang fields in the year of 2015 and 2016, which explained from 3.74 to 11.89% of the total phenotypic variation (PV) with LOD scores ranging 2.50 to 6.96. They were located on 20 chromosomes except Chr04, Chr08, Chr13, Chr16, Chr18 and Chr25. Among them, 35 QTLs (66%) had negative additive effects, indicating that their favorable alleles come from *G. barbadense*, which enhanced VW resistance and decremented DI by 2.64 to 13.23. On the other hand, 18 QTLs (34%) had positive additive effects, indicating that the *G. barbadense* alleles decremented VW resistance and enhanced phenotypic DI values by 2.27 to 19.47. Thirty-one QTLs were identified in 2015 and 86 QTLs in 2016, of which eleven ones were found in the both years. The highest number of QTLs (11) was detected on Chromosome 5 (Figure 3, Table S1).

**Figure 3.**
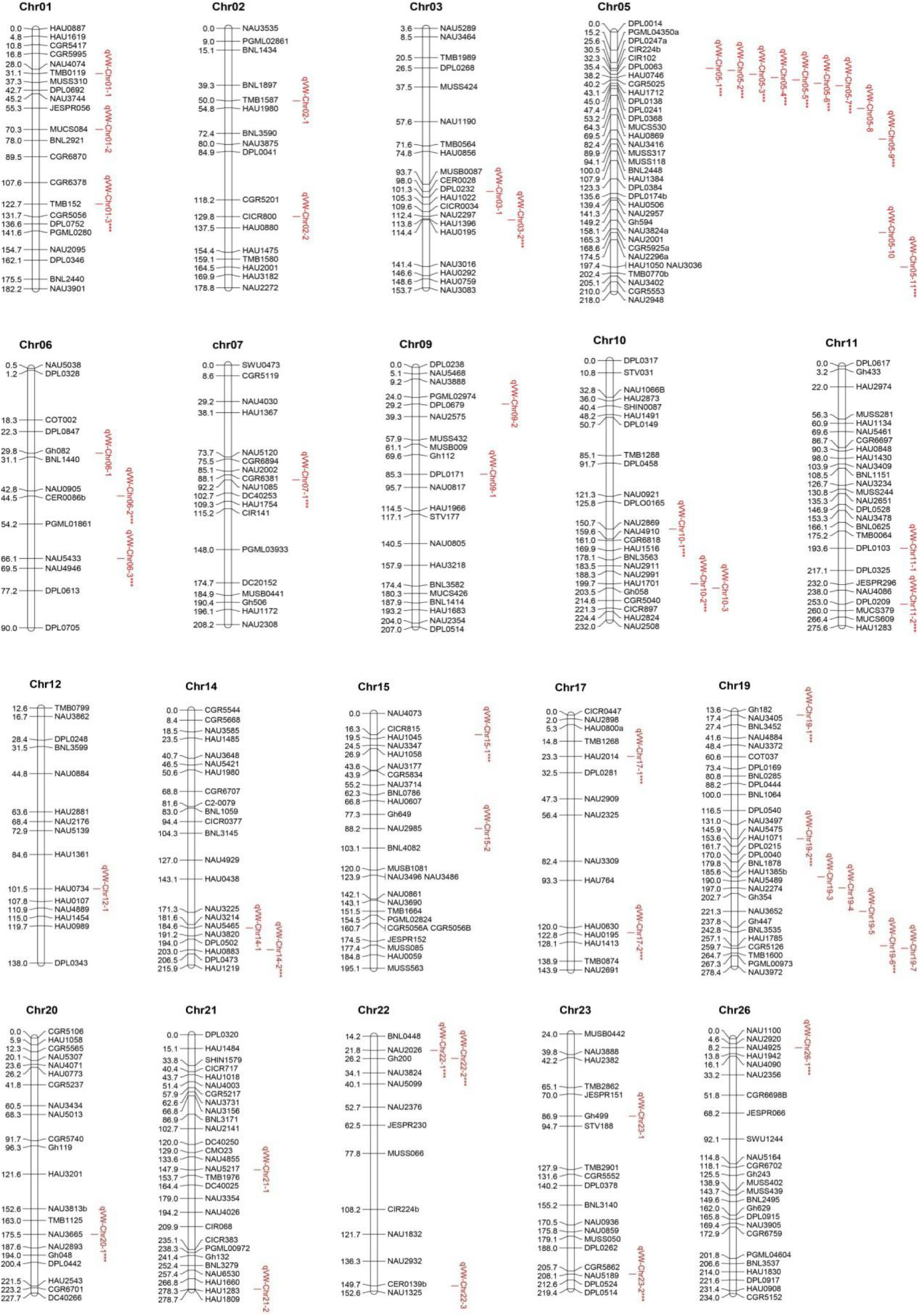
Identification of QTLs for VW disease index and linkage map in BC5F3:5 populations. Note: stars indicate stable QTLs

#### 2.4.1 QTLs for VW resistance in Anyang in 2015

In July 2015, there were ten QTLs identified in Anyang and mapped on 5 chromosomes, explaining 4.39–11.89% of overall PV with LOD scores ranging 2.89–4.87, of which five ones were found on Chr05 while two ones on Chr19. All QTLs except *qVW-Chr05-8* and *qVW-Chr19-5* had negative additive effects, indicating that their favorable alleles derived from donor parent Hai1 incremented VW resistance and decremented phenotypic DI by 4.75-7.98 (Table S1).

In August 2015, thirteen QTLs were identified at Anyang and mapped on 8 chromosomes, explaining 3.79–7.67% of the overall phenotypic variation with LOD scores ranging 2.51–5.22. Five QTLs were found on Chr05 and two QTLs on Chr19, which was consistent with the results in July 2015. Except for *qVW-Chr01-1*, *qVW-Chr12-1* and *qVW-Chr26-1*, the whole QTLs had negative additive effects, which suggested that donor parent *G. barbadense* alleles incremented VW resistance and decremented DI by 2.64-13.23 (Table S1).

#### 2.4.2 QTLs for VW resistance at Xinjiang in 2015

In July 2015, there were six QTLs detected at Xinjiang, which were mapped on 6 Chromosomes with 3.78–9.33% of the total PV explained. All the QTLs showed positive additives, which suggested the Hai1 alleles decremented resistance against VW and incremented phenotypic DI by 2.27-13.25 (Table S1).

In August 2015, two QTLs were found at Xinjiang, namely as *qVW-Chr05-10* and *qVW-Chr06-1* which were mapped on Chr5 and Chr6 with 5.00 and 5.59% of PV and LOD scores of 3.35 and 3.94, respectively. These QTLs also presented positive additives, suggesting their alleles derived from *G. barbadense* decreased resistance of the disease and increased DI by 2.81 and 9.40 (Table S1).

#### 2.4.3 QTLs for VW resistance in Anyang in 2016

In July 2016, there were fourteen QTLs detected at Anyang and mapped on 9 chromosomes, explaining 4.27–7.71% of the total PV. Four QTLs were located on Chr05, while each two QTLs were identified on Chr06 and Chr19, respectively. All the QTLs had negative additives, which suggested their parent Hai alleles incremented VW resistance and decremented DI by 4.43-9.57 (Table S1).

In August 2016, ten QTLs were recorded at Anyang and mapped on 8 chromosomes, explaining 3.76–6.15% of the overall PV, of which three ones were identified on Chr05. All the QTLs except *qVW-Chr02-3* had negative additives, suggesting their alleles derived from parent Hai1 incremented resistance and decremented DI by 4.19-10.47 (Table S1).

#### 2.4.4 QTLs for VW resistance in Xinjiang in 2016

In July 2016, there were twenty-eight QTLs detected at Xinjiang and mapped on 14 chromosomes with 3.74–11.14% of total PV explained, of which LOD score ranging was 2.55–6.96. In addition, nine QTLs were found on Chr05, and three QTLs were located on Chr19. All the QTLs except *qVW-Chr09-1*, *qVW-Chr10-2*, *qVW-Chr15-1*, and *qVW-Chr22-2* had negative additives, suggesting their parent Hai1 alleles enhanced resistance against VW and decreased DI by 2.96-7.65 (Table S1).

In August 2016, thirty-four QTLs were found at Xinjiang and mapped on 15 chromosomes, explaining 3.79–10.22% of total PV. Nine QTLs were identified on Chr05, while five and three QTLs were separately located on Chr19 and Chr10. Except for *qVW-Chr10-2*, *qVW-Chr10-3*, *qVW-Chr15-1*, and *qVW-Chr22-2*, all the QTLs had negative additives, which suggested their alleles derived from parent *G. barbadense* incremented resistance against VW and decremented phenotypic value of DI by 3.94-10.48 (Table S1).

### 2.5 Identification of stable QTLs over environments and developmental periods

In total, 53 QTLs of VW disease index were detected in CSSLs during different stages of growth and environments, which were separately located on 20 different chromosomes. There were 11 and 7 QTLs identified on Chr05 and Chr19, respectively, and each 3 QTLs were separately located on Chr01, Chr06, Chr10, and Chr22. Each 2 QTLs were found on Chr02, Chr03, Chr09, Chr11, Chr14, Chr15, Chr17, Chr21, and Chr23, respectively, while Chr07, Chr12, Chr20, Chr24, and Chr26 separately contained only 1 QTL (Table S1).

Among 53 QTLs, 29 stable QTLs were identified in at least two environment, explaining 3.74-11.89% of the overall PV (Table 4). There were 25 stable QTLs (86%) showing negative additive effects, which suggested thier Hai1 alleles enhanced resistance against VW and decreased phenotypic DI. Among 29 stable QTLs, Chr05 harbored 09 stable QTLs, and Chr19 contained 3 stable QTLs. Each 2 stable QTLs were separately located on Chr06, Chr10, Chr17, and Chr22, while Chr01, Chr03, Chr07, Chr11, Chr14, Chr15, Chr20, Chr23, and Chr26 contained 1 stable QTL, respectively.

**Table 4.**
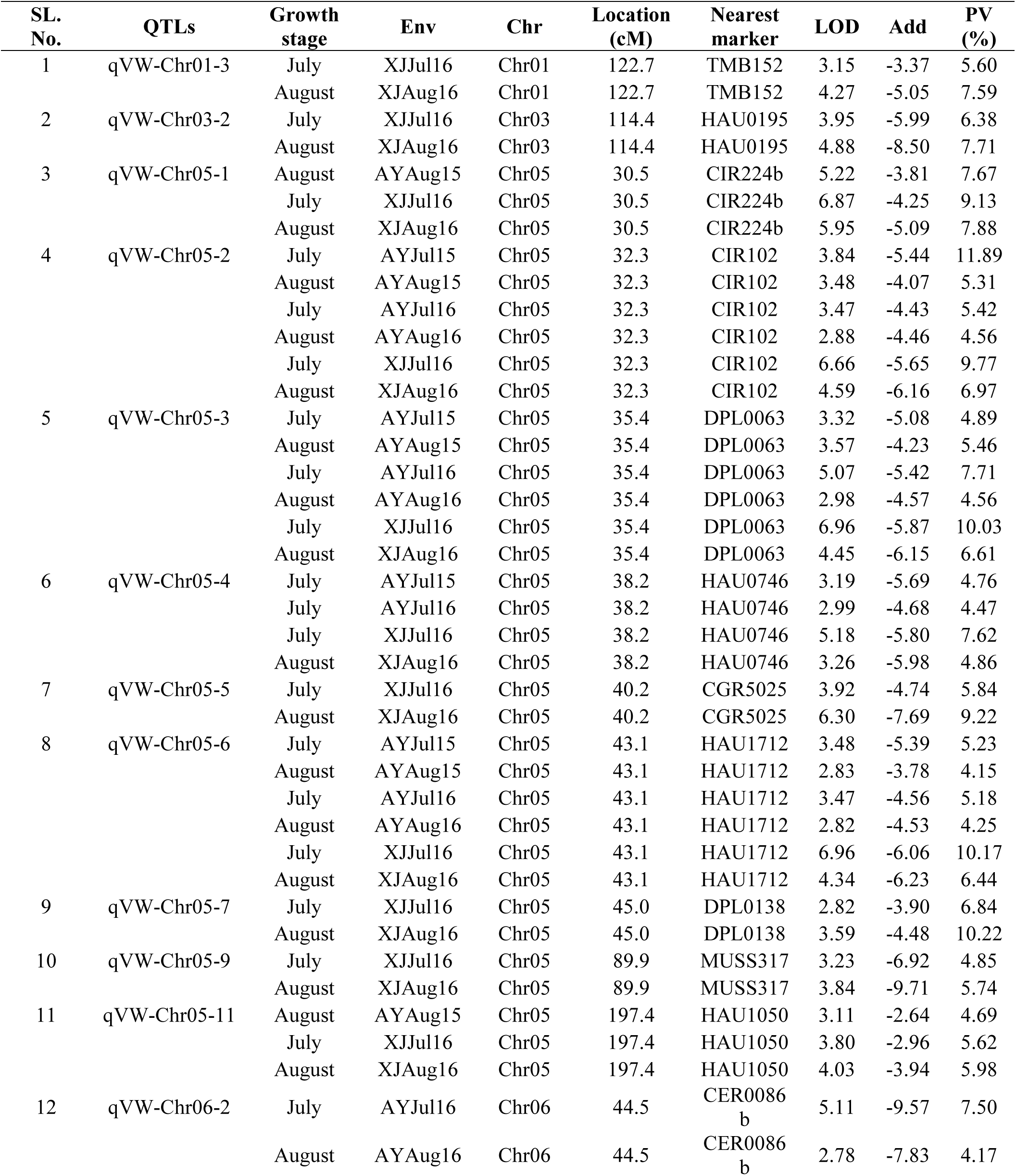
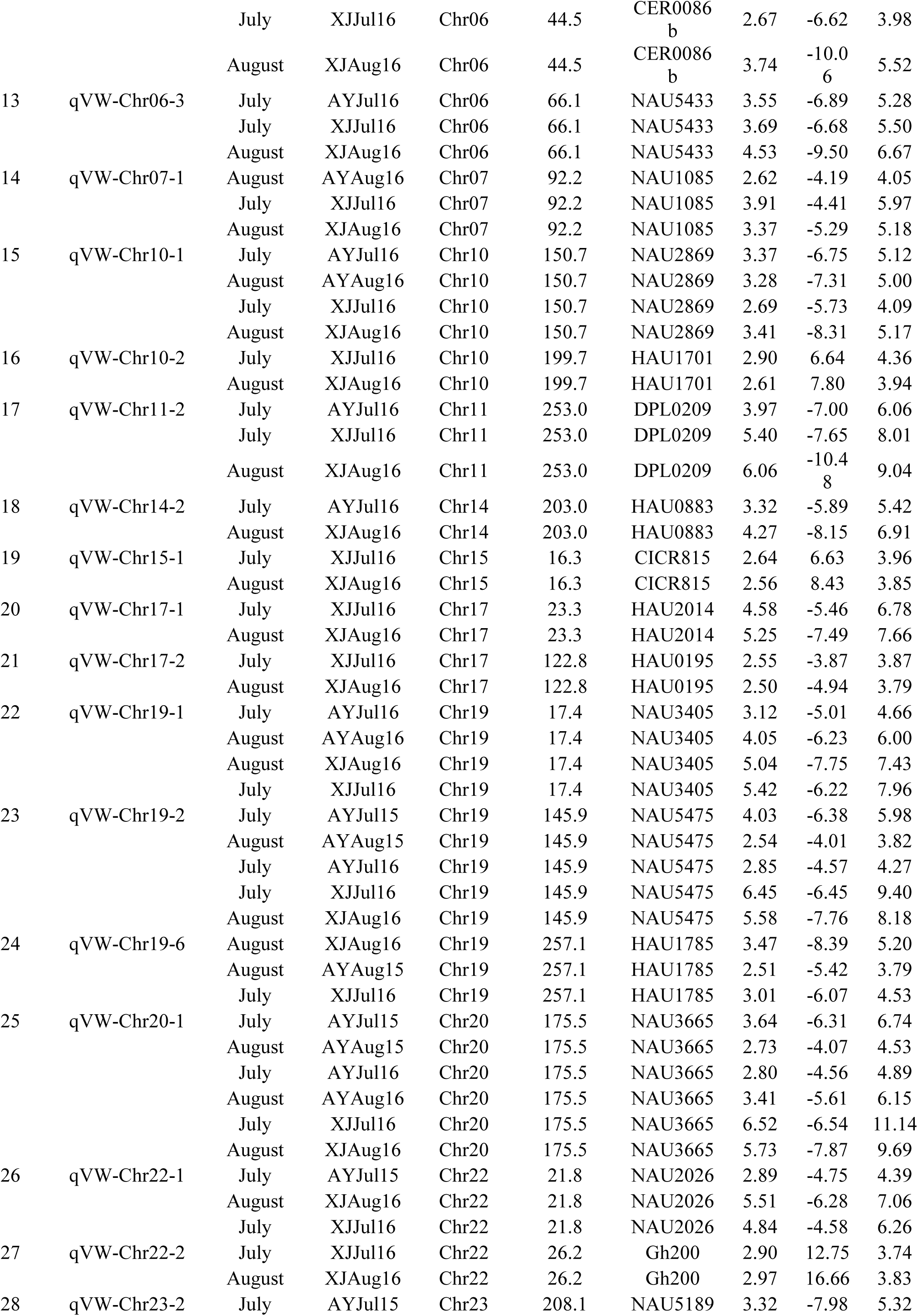
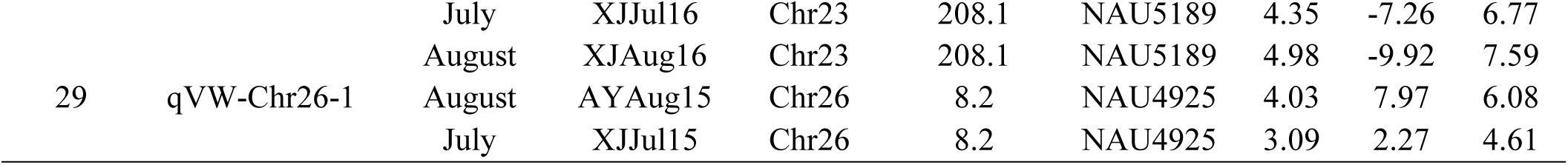
Identification of QTLs for VW disease index during different development and environments in BC5F3:5 populations

Four stable QTLs, namely as *qVW-Chr05-2*, *qVW-Chr05-3*, *qVW-Chr05-6*, and *qVW-Chr20-1*, were detected in six environments explaining 4.56-11.89%, 4.56-10.03%, 4.15-10.17% and 4.53-11.14% of PV, respectively. Only one stable QTL (*qVW-Chr19-2*) was identified in five environments with 3.82-9.40% of the observed PV, while three stable QTLs (*qVW-Chr05-4*, *qVW-Chr10-1*, and *qVW-Chr19-1*) were investigated in four environments separately explaining the observed PV of 4.47-7.62%, 4.09-5.17%, and 4.66-7.96%. Moreover, there were nine stable QTLs detected in three environments, namely as *qVW-Chr05-1*, *qVW-Chr05-11*, *qVW-Chr06-2*, *qVW-Chr06-3*, *qVW-Chr07-1*, *qVW-Chr11-2*, *qVW-Chr19-6*, *qVW-Chr22-1*, and *qVW-Chr23-2*, which presented 7.67-9.13%, 4.69-5.98%, 3.98-5.52%, 5.28-6.67%, 4.05-5.97%, 6.06-9.04%, 3.79-5.20%, 4.39-7.06%, and 5.32-7.59% of the observed PV, respectively. Twelve stable QTLs were detected in two environments with overall 3.74-10.22% of PV. The stable QTLs, including *qVW-Chr05-2*, *qVW-Chr05-3*, *qVW-Chr05-6*, *qVW-Chr05-7*, and *qVW-Chr20-1*, had major effects and explained 11.89%, 10.03%, 10.17%, 10.22% and 11.14% of the observed PV, respectively (Table 4).

### 2.6 QTL hotspots and meta-analysis

Based on Meta-analysis, 32 QTL hotspot regions were totally detected on 18 chromosomes, including Chr01, Chr03, Chr05, Chr06, Chr07, Chr09, Chr11, Chr12, Chr14, Chr15, Chr17, Chr19, Chr20, Chr21, Chr22, Chr23, Chr24 and Chr26 (Figure S1, Table 5). Among them, 17 QTL hotspot regions were consistent with those detected earlier by [7, 22, 33] (Table 5), and the other 15 were identified as novel ones. Each 3 QTL hotspot regions were separately located on Chr05, Chr19, and Chr26, while each 2 QTL hotspot regions were detected on Chr01, Chr03, Chr07, Chr09, Chr20, Chr21, Chr22, and Chr23, respectively. In addition, Chr06, Chr11, Chr12, Chr14, Chr15, Chr17, and Chr24 separately contained 1 QTL hotspot region (Table 5).

**Table 5.** QTL hotspots detected for VW resistance on the consensus map through meta-analysis

| Hotspot name | Chr | Location (cM) | No. of QTLs | No. of QTLs in this paper | Reported earlier |
| --- | --- | --- | --- | --- | --- |
| Chr01-DI-Hotspot-1 | Chr01 | 14-33 cM | 6 | 3 |  |
| Chr01-DI-Hotspot-2 | Chr01 | 34-55 cM | 5 | 0 |  |
| Chr03-DI-Hotspot-1 | Chr03 | 20-34 cM | 5 | 1 | Shi et al., 2016 |
| Chr03-DI-Hotspot-2 | Chr03 | 34-44 cM | 5 | 1 |  |
| Chr05-DI-Hotspot-1 | Chr05 | 25-36 cM | 18 | 8 | Shi et al., 2016 |
| Chr05-DI-Hotspot-2 | Chr05 | 31-42 cM | 5 | 2 | Shi et al., 2016;<br>Said et al., 2015 |
| Chr05-DI-Hotspot-3 | Chr05 | 39-54 cM | 17 | 1 | Shi et al., 2016;<br>Zhang et al., 2015 |
| Chr06-DI-Hotspot-1 | Chr06 | 35-51 cM | 6 | 2 |  |
| Chr07-DI-Hotspot-1 | Chr07 | 51-76 cM | 6 | 0 | Zhang et al., 2015 |
| Chr07-DI-Hotspot-2 | Chr07 | 178-193 cM | 8 | 1 |  |
| Chr09-DI-Hotspot-1 | Chr09 | 34-60 cM | 9 | 2 | Zhang et al., 2015 |
| Chr09-DI-Hotspot-2 | Chr09 | 87-93 cM | 2 | 0 |  |
| Chr11-DI-Hotspot-1 | Chr11 | 72-99 cM | 6 | 2 |  |
| Chr12-DI-Hotspot-1 | Chr12 | 13-28 cM | 6 | 1 | Shi et al., 2016 |
| Chr14-DI-Hotspot-1 | Chr14 | 19-34 cM | 7 | 2 | Shi et al., 2016 |
| Chr15-DI-Hotspot-1 | Chr15 | 41-68 cM | 10 | 1 |  |
| Chr17-DI-Hotspot-1 | Chr17 | 6-23 cM | 4 | 2 |  |
| Chr19-DI-Hotspot-1 | Chr19 | 2-27 cM | 5 | 1 | Shi et al., 2016;<br>Zhang et al., 2015 |
| Chr19-DI-Hotspot-2 | Chr19 | 32-55 cM | 4 | 0 | Zhang et al., 2015 |
| Chr19-DI-Hotspot-3 | Chr19 | 123-148 cM | 7 | 4 | Shi et al., 2016 |
| Chr20-DI-Hotspot-1 | Chr20 | 12-24 cM | 4 | 0 | Shi et al., 2016;<br>Zhang et al., 2015 |
| Chr20-DI-Hotspot-2 | Chr20 | 26-45 cM | 5 | 1 |  |
| Chr21-DI-Hotspot-1 | Chr21 | 3-29 cM | 6 | 0 | Zhang et al., 2015 |
| Chr21-DI-Hotspot-2 | Chr21 | 35-60 cM | 7 | 2 |  |
| Chr22-DI-Hotspot-1 | Chr22 | 0-25 cM | 5 | 1 | Zhang et al., 2015 |
| Chr22-DI-Hotspot-2 | Chr22 | 30-54 cM | 7 | 2 |  |
| Chr23-DI-Hotspot-1 | Chr23 | 40- 65 cM | 30 | 1 | Zhang et al., 2015 |
| Chr23-DI-Hotspot-2 | Chr23 | 67-92 cM | 26 | 1 |  |
| Chr24-DI-Hotspot-1 | Chr24 | 0-25 cM | 5 | 1 | Zhang et al., 2015 |
| Chr26-DI-Hotspot-1 | Chr26 | 4-29 cM | 6 | 1 | Zhang et al., 2015 |
| Chr26-DI-Hotspot-2 | Chr26 | 33-54 cM | 5 | 0 |  |
| Chr26-DI-Hotspot-3 | Chr26 | 85-102 cM | 5 | 0 |  |

Among 32 QTL hotspot regions, 9 hotspot regions located on seven different chromosomes had more QTLs (Figure S1, Table 5), which could be very important for further studies and utilized for molecular breeding via MAS. As for chr05, 40 QTLs were selected to project on consensus chromosome 05 (Cons.Chr05), resulting in 3 identified QTL hotspot regions. There were 18, 5, and 17 QTLs on Chr05-DI-Hotspot-1, Chr05-DI-Hotspot-2, and Chr05-DI-Hotspot-3, respectively (Figure 4, Table 5). Eleven QTLs were selected to project on chromosome 09 (Cons.Chr09), and 2 QTL hotspot regions were identified, of which Chr09-DI-Hotspot-1 had 9 QTLs, while Chr09-DI-Hotspot-2 had 2 QTLs. Sixteen QTLs were identified and projected on consensus Chr19 to perform meta-analysis, identifying 3 QTL hotspot regions. Chr19-DI-Hotspot-1, Chr19-DI-Hotspot-2 and Chr19-DI-Hotspot-3 contained 5, 4 and 7 QTLs, respectively. Twelve QTLs were selected to project on chromosome 22 (Cons.Chr22), and 2 QTL hotspot regions were identified, of which Chr22-DI-Hotspot-1 had 5 QTLs, while Chr22-DI-Hotspot-2 had 7 QTLs (Figure 4). Fifty six QTLs were selected to project on Cons.Chr23, identifying 2 QTL hotspot regions. Chr23-DI-Hotspot-1 and Chr23-DI-Hotspot-2 contained 30 and 26 QTLs, respectively. Sixteen QTLs were selected to project on Cons.Chr26, and 3 QTL hotspot regions were identified. Chr26-DI-Hotspot-1, Chr26-DI-Hotspot-2 and Chr26-DI-Hotspot-3 contained 6, 5 and 5 QTLs, respectively. The details of all QTLs are described in Table 5.

**Figure 4.**
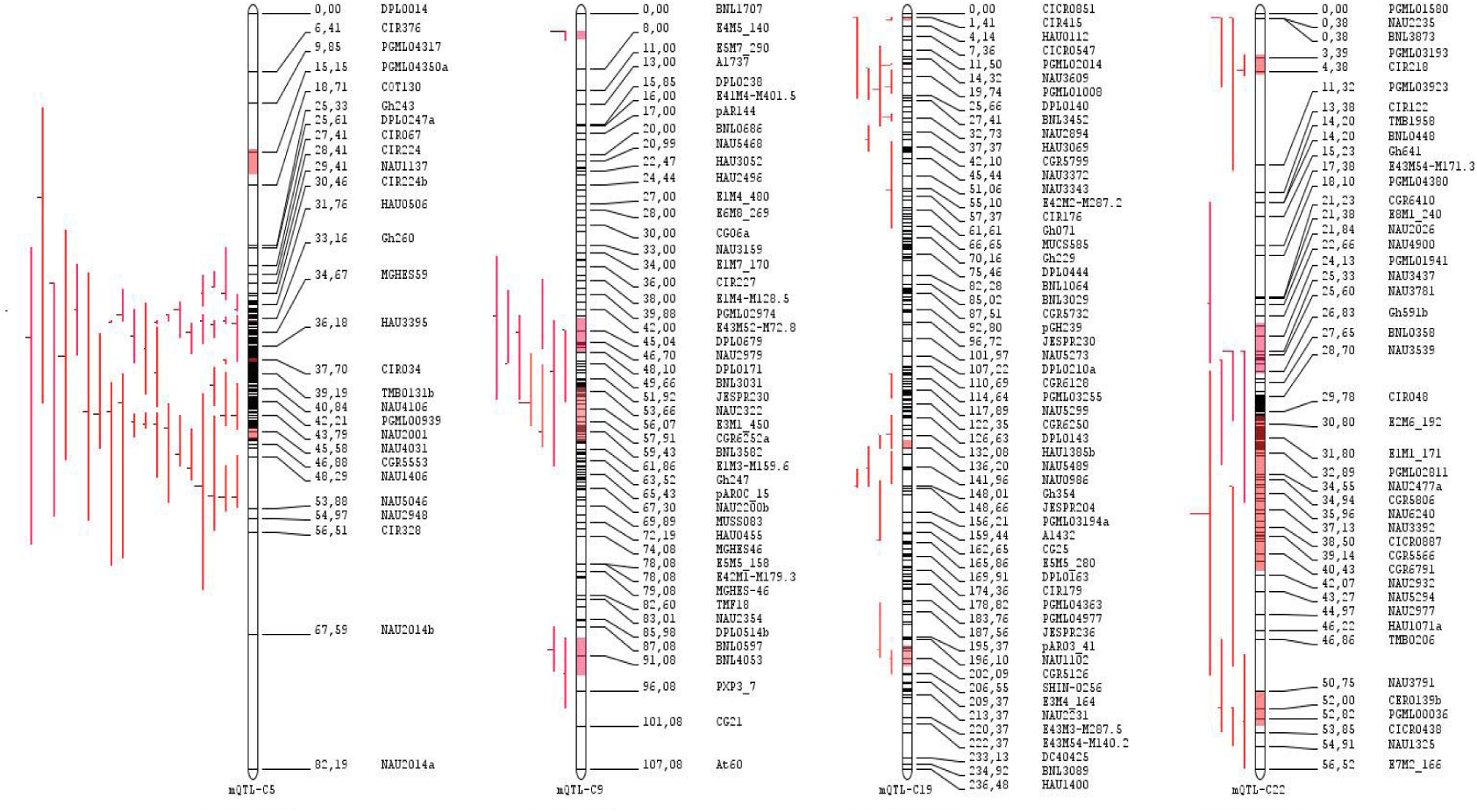
QTL hotspots and QTLs for VW resistance on the consensus map by a meta-analysis. Consensus Chromosome 05 (Cons.Chr05) has three hotspots, Cons.Chr09 has 2, Cons.Chr19 has three and Cons.Chr22 has 2 hotspots.

As for the hotspots on Cons.Chr05, Chr05-DI-Hotspot-1 from the 25–36 cM region was located between markers Gh243 and HAU3395, and Chr05-DI-Hotspot-2 from 31 to 42 cM region and Chr05-DI-Hotspot-3 from 39 to 54 cM region were separately located between markers NAU3204 and CIR301 and between markers TMB0131b and NAU2948. There were two hotspots on Cons.Chr09, and Chr09-DI-Hotspot-1from the 34–60 cM region and Chr09-DI-Hotspot-2 from 87 to 93 cM region were located between markers CGR6170 and CGR6719 and between markers BNL0597 and BNL4053, respectively. With regard to the hotspots on Cons.Chr19, Chr19-DI-Hotspot-1 from the 2–27 cM region was located between markers CIR415 and BNL3452, and Chr19-DI-Hotspot-2 from 32–55 cM region and Chr19-DI-Hotspot-3 from 123 to 148 cM region were separately located between markers NAU2894 and COT037 and between markers DPL0216 and Gh354. Moreover, three hotspots were identified on Cons.Chr26, of which Chr26-DI-Hotspot-1 from 4 to 29 cM region was located between markers HAU1845 and DPL0888, while Chr26-DI-Hotspot-2 from 33 to 54 cM region and Chr26-DI-Hotspot-3 from 85 to 102 cM region were lacated between markers NAU2356 and CIR167 and between markers C2-0528 and DPL1283, respectively.

## 3. Discussion

### 3.1 Field status and phenotypic assessment

Without the inoculation provision and just under natural environmental conditions, a population of CSSLs developed from interspecific cross between Upland cotton CCRI36 and Sea Island cotton Hai1, which has been investigated for resistance against VW together with parents and controls. The VW resistance was assessed based on the leaf tissue damage in the mature stages, of which the results indicated the parent Hai1 appeared to be more resistant to the disease compared to CCRI36, while the control Jimian11 displayed slightly higher susceptibility over CCRI36. Most of the CSSLs exhibited higher DI values than mid parents (Table 2), and this unclear phenomenon might be due to DI values fluctuation across the environments. The same remark was made in a study using an interspecific chromosome segment line with different VW strains and according to the authors, that fact can be explained the resistance to different VW isolates is controlled by distinct single genes and that in the presence of a mixture of isolates, interactions occurred [19].

Over different years of study and across variable environments, the investigated population of CSSLs has displayed a broad range of sensitivity ranging between highly susceptible to highly resistant. Having taken the previous studies [40] into consideration, the hypothesis came into being regarding inheritance of VW in recessive fashion, which is that both the paternal and maternal contributors should harbor genetic factors for resistance. For the verification of the generated hypothesis, the CSSL population has been investigated on phenological basis over different environments at various growth stages. In this study, we observed that DI values susecptible to VW infection were higher in August than those in July, to be specific to presenting that the susceptible control (Jimian11) showed above 35% DI values except in XJJul15 and AYJul16, while the DI values of CCRI36 were lower than 35% except in AYAug15 and XJAug16 (Table 2). This lesser DI percentage is the evidence for the occurrence of high pressure projected by variable VW strains under natural environmental conditions. Few more reasons behind this phenological variation include intensity and virulence of strains, fungal amount in soil and developmental stages as well as environmental influences [41]. The similar findings have been reported earlier in which the host plant proved to be resistant against inoculum of VW while remained susceptible under natural environmental conditions [21]. We also have synchrony with previous findings with a display of lesser disease index (DI<40%) by CCRI36 progenitor whereas some of the offspring depicted a prominent resistance level comparable to susceptible control Jimian11. Besides this, a noteworthy level of transgressive segregation has been witnessed under field conditions, which are in accordance with previous reports [42, 43]. Across different environment during whole investigation period, few CSSLs remained consistent in resistance display to pressurizing mixture of strains present in the vicinity as compared to most of the lines which displayed a high level of susceptibility (Figure 2). This fact can be justified by the presence of wider range of environmental variation occurrence during two experimental years of study, where the VW strains keep on changing their genetic make up for being more resistant. Previous reports [19] justified our such findings for the confirmation of reality that there must exists an antagonistic interaction between resistance QTLs/genes and different strains of fungi plus large number of genes are responsible for controlling the resistance mechanism against *V*. *dahliae* isolates.

The phenological parameters measured in two years of study at both locations depicted rare weak correlations. Expression of different genetic factors in variable environments at different growth stages confirmed the reason behind weak correlation coefficient values (Table 3). It realizes the fact regarding alteration of genes on exposure to VW strains at varying growth stages. In a study on backcross inbreed lines regarding VW resistance, there observed a weak but positive correlation among disease index under field conditions [44].

Due to varying environmental stresses in both years at two locations, erroneous frequency was very high and because of this heritability values ranged between weak to moderate only. This happening suggests a wider range of phenology regarding DI has been caused by varying environmental influences. However, this is not a surprising truth as cotton resistance levels to *V*. *dahliae* are greatly inclined to environmental influences, resistance genes, inoculum concentrations and their interactions [45].

### 3.2 Genetic Map used for QTLs identification

Through utilization of hybridization technique including interspecific [7, 18, 21, 42, 43, 45–48] and intraspecific [21, 46, 47, 49, 50] crossing wide range of genetic maps have been constructed. However, lesser genome coverage i.e. < 50% has been achieved by using interspecific crossing, which appeared as bottleneck in the detection of QTLs from whole genome with ultra-resolution. The fact has been proved by the discovery of about 57.90% of tetraploid cotton genome from Zhang et al. [7] study, 27% i.e. 1143.1cM and 35% with 279 markers of genome coverage in Fang et al. [21] and [47] reports. To date, one exclusive report has found that covered more than 50% of genome i.e. 55.7% accounting for 882 genetic markers in total, including 414 SNPs, 36 RGA-RFLPs (resistance gene analog-amplified fragment length polymorphism) and 432 SSRs. Therefore, the whole genome coverage of allotetraploid cotton with resistant QTLs for VW is not yet to be achieved. This study paced to cover approximately 100% of cotton genome enclosing about 5115.6cM [37], which is really a comprehensive distance accomplished so far. It’s neonatal to take in account all the 26 genetic threads of allotetraploid cotton with use of CSSLs in quest of QTLs for VW resistance. An announce-worthy amount of QTLs (53) were identified to be related to VW resistance from 20 chromosomes, which exposed the reality that these QTLs are extensively distributed in whole genome chromosomes. These results would be not easy to achieve if *G.barbadense* genome will be used as template with restricted amount of markers and lesser polymorphism.

### 3.3 Distribution of QTLs of Verticillium wilt through the whole genome

There were fewer chromosomes yet to have been explored regarding VW resistance QTLs in the previous studies, specifically including Chromosome 6, Chromosome 10, Chromosome 12, and Chromosome 18 together with almost 100 plus related QTLs [45, 51], which left these gaps from completing the whole tetraploid genome. Our findings have contributed plenty of valuable information to filling up there gaps to greater extent, leaving just Chromosome 18 to be explored. There were three QTLs detected on Chromosome 6 and 10, while only one DI QTL was identified on Chromosome 12. Like previous findings such as Zhang et al. [7] from meta-analysis done by different researchers, we also remained unable to discover any hotspot region on Chromosomes 10 and 18. However, few chromosomes were found to be heavily loaded with DI QTLs like Chromosome 5 with 11 DI QTLs, and Chromosome 19 with 7 DI QTLs. Each 3 QTLs were separately located on Chromosome 1 and 22 like Chromosome 6 and 10 as mentioned earlier. Also in our findings we remained, successful in identifying some stable QTLs across six different environments, which was not the case in any of the previous reports.

As mentioned earlier, 20 chromosomes were explored in our study with 53 QTLs using BC_5_F_3:5_ populations, of which 30 QTLs were located on A sub-genome chromosomes covering Chr01, Chr02, Chr03, Chr05, Chr06, Chr07, Chr09, Chr10, Chr11 and Chr12 accounting 56.66%, while 23 QTLs were explored on D sub-genome covering Chr14, Chr15, Chr17, Chr19, Chr20, Chr21, Chr22, Chr23, Chr24 and Chr26 estimating about 43.44%. There results provided an evidence of the fact that A sub-genome enclosed more resistant QTLs for VW resistance as compared to D sub-genome. Consistent discoveries have been made by Yang et al. [46], Ning et al. [47] and Bolek et al. [42].

### 3.4 Stability with earlier studies VW resistance QTLs

In this study, 53 QTLs related to VW resistance were totally identified in 300 CSSLs. Among all the QTLs, 35 ones (66%) had negative additive effects, which indicated that the *G. barbadense* alleles increased *Verticillium* wilt resistance and decreased disease index values by about 2.64 to 13.23. On the other hand, 18 QTLs (34%) had positive additives effects, which indicated that the *G. hirsutum* alleles enhanced VW wilt resistance and decremented phenotypic disease index values by about 2.27 to 19.47. As for different years, 31 QTLs were identified in the year of 2015, while 86 QTLs in the year of 2016, of which 11 QTLs were found in the both years. The maximum number of QTLs (11) was detected on Chr05 (Figure 3, Table S2).

Among 53 QTLs, 29 QTLs were detected consistently in at least two environments, which were deemed as stable QTLs. Out of 29 stable QTLs, 25 QTLs (86%) had negative additive effects, which indicated that the *G. barbadense* alleles incremented VW resistance and decreased DI. Based on Meta-analysis of the identified 53 QTLs, 10 QTLs were consistent to previously identified QTLs, and they had common SSR markers [19, 45–47, 52]. One QTL, *qVW-Chr01-3* positioned on Chr01 for VW resistance was the similar as Ning’s *qVW-A1-1* [47], which were identified with common markers of Gh215. Another QTL, *qVW-Chr03-2* was the similar as *qVW-C3-2* in the results of Shi et al. [22], and they were associated with the shared marker CER0028. In addition, *qVW-Chr05-1* on Chr05 was similar as Shi et al’s *qVW-C5-1* [22] based on common marker CIR224b. The *qVW-Chr05-11* mapped on Chr05 was similar as the *qVLBP2-A5-1RIL* in the results of Yang et al. [46], which were associated with shared markers NAU5210. The QTL *qVW-Chr05-4* was similar as the *qVW-C5-3* in the results of Shi et al [22] with the association of shared marker HAU0746 [22]. The *qVW-Chr07-1* was similar as *qVW-A7-1* in the results of Ning et al., [47] based on shared marker Gh527. *qVW-Chr09-1* mapped on Chr09 was the similar as Shi’s *qVW-C9-1* [22], with the association of common markers of DPL0783. The QTL *qVW-Chr12-1* was the similar as *qVWR-06-C12* in the results of Zhang et al. [7], which were associated with the common marker CIR272. Besides, *qVW-Chr23-2* was similar as Fang’s *qDR52T2-C23-2* [48] associated with the shared marker DPL1938. Lastly, the QTL *qVW-Chr05-1* was similar as the *qVW-C5-2* in the results of Shi et al [22] with the association of shared marker CIR102 [22]. The remaining 43 QTLs for VW resistance could be allowed as novel ones in this study.

Based on meta-analysis, 32 QTLs hotspot regions were detected, of which 17 ones were consistent with the earlier studies [7, 22, 33], while another 15 ones were novel and unreported hotspot regions (Figure 4, Table 5). These hotspot regions and QTLs could be very important information for further comparative studies and utilized for marker assisted selection.

### 3.5 Further utilization of QTLs for VW resistance

According to previous reports on the CSSLs in cotton, the prominent characteristics of high fiber quality and high yielding traits have deliberately been explained [53–58]. Nowadays in this whole experimental study, a total of 300 CSSLs from Upland cotton CCRI36 and Sea Island cotton Hai1 have been keenly investigated regarding their resistance to VW. The segments of chromosome introgressed from *G*. *barbadense* into *G*. *hirsutum* made these lines little bit different from their recurrent parent by reducing the influences of genetic background of recipient, which makes the CSSLs as efficient breeding materials to conduct quantitative genetics researchs. Thus the experimented work proves to be beneficial in paving the way towards whole genome study of cotton by laying a solid platform stuffed with molecular findings related to fine mapping, functional genomics, gene pyramiding and ultimately marker assisted breeding.

## 4. Materials and Methods

### 4.1 Plant materials and development of cotton CSSLs

Mapping population based on 300 CSSLs along with their parents, specifically as CCRI36 (*G. hirsutum*) as recurrent while Hail (*G. barbadense*) as donor parent, was sown at the farm area of ICR, CAAS (Anyang, Henan) and Shihezi, Xinjiang Province, respectively. The reason behind selection of Hail as donor parent is its characteristic features of producing high quality fiber, resistant genes residence for VW in its genome and also the presence of glandless producing factors which act in dominant fashion [59]. However, CCRI36 developed by ICR, CAAS (State Approval Certificate of Cotton 990007) [36] is a commercially grown renowned variety of upland cotton has the obvious property of high yielding as well as early maturing in growth patterns but susceptible to *Verticillium* wilt. The two cultivars Hail and CCRI36 used as paternal and maternal parents were hybridized followed by backcross in 2003 at Anyang to construct CSSLs. In 2009, a mapping population comprising 2660 plants of BC_5_F_3_ was obtained by using CCRI36 as recurrent parent. In 2010 and 2011, BC_5_F_3:4_ population was planted via plant-to row method at Anyang and Xinjiang, respectively. In 2014, at Xinjiang province, BC_5_F_3:5_ population was grown again. From these populations, a random selection process was conducted and 300 CSSLs were obtained for the evaluation of VW disease index. These selected lines were then grown at Anyang and Xinjiang in 2015 and 2016, respectively. The details of development of CSSLs was brought about by following the same procedure as described earlier [60]. Stable performance regarding resistance to VW was displayed by some lines in multiple environments over different years of study.

### 4.2 Field investigations and experimental design

Two field stations of ICR, CAAS in Anyang, Henan and Shihezi, Xinjiang were used to grow the experimental material for two years. In 2015 and 2016, phenotypic data were collected in months of July and August from Anyang and Xinjiang, respectively. Under natural environmental conditions, there occurred intensive attack of *V. dahliae* strains. Randomized complete block design (RCBD) under two replications was established for study. By following the specifications prescribed for crop management according to the locality, seeds were sown in single row plots. At research farm areas of Anyang, planting rows were kept 5 m long with an interval of 0.8 m whereas thinning of seedlings was done upto 20 plants in a row. However, in Xinjiang row length was kept at 3 m with plant to plant distance of 0.1 m following two-narrow by row plots methodology. Row spacing alternation was 0.1 m by 0.66 m. The detail of field layout is mentioned in Table 6. Wide/narrow row to row distance pattern was followed and plastic membranes were utilized for covering of seedlings. Standard agronomic performs were established during whole experiment at all locations.

**Table 6.**
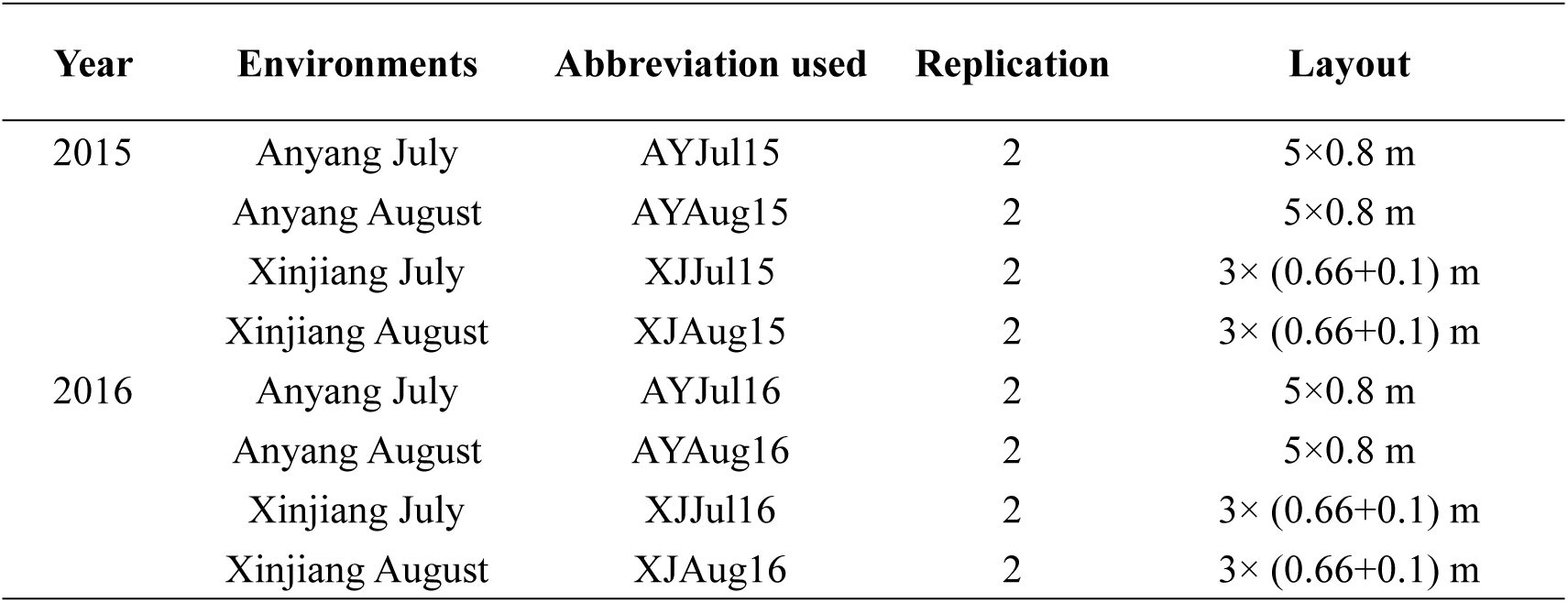
Details of 8 environments of fields used to evaluate CSSL population

| Year | Environments | Abbreviation used | Replication | Layout |
| --- | --- | --- | --- | --- |
| 2015 | Anyang July | AYJul15 | 2 | 5×0.8 m |
|  | Anyang August | AYAUG15 | 2 | 5×0.8 m |
|  | Xinjiang July | XJJul15 | 2 | 3× (0.66+0.1) m |
|  | Xinjiang August | XJAUG15 | 2 | 3× (0.66+0.1) m |
| 2016 | Anyang July | AYJul16 | 2 | 5×0.8 m |
|  | Anyang August | AYAUG16 | 2 | 5×0.8 m |
|  | Xinjiang July | XJJul16 | 2 | 3× (0.66+0.1) m |
|  | Xinjiang August | XJAUG16 | 2 | 3× (0.66+0.1) m |

### 4.3 Verticillium wilt phenotypic evaluation

For scoring of diseased portion of plant, a percentage based scale was used for evaluation ranging between 0-4 [61]. The scale used is a standard one being used deliberately in China especially for *Verticillium* disease rating indices by classifying the damaged portion of matured stage leaves into five groups [46, 51, 62]. The scoring pattern is considered in ascending order regarding resistance level accounting 0-2 as resistant and 3-4 as susceptible. The disease rating scale of VW is comprehensively discussed in Table 7.

**Table 7.**
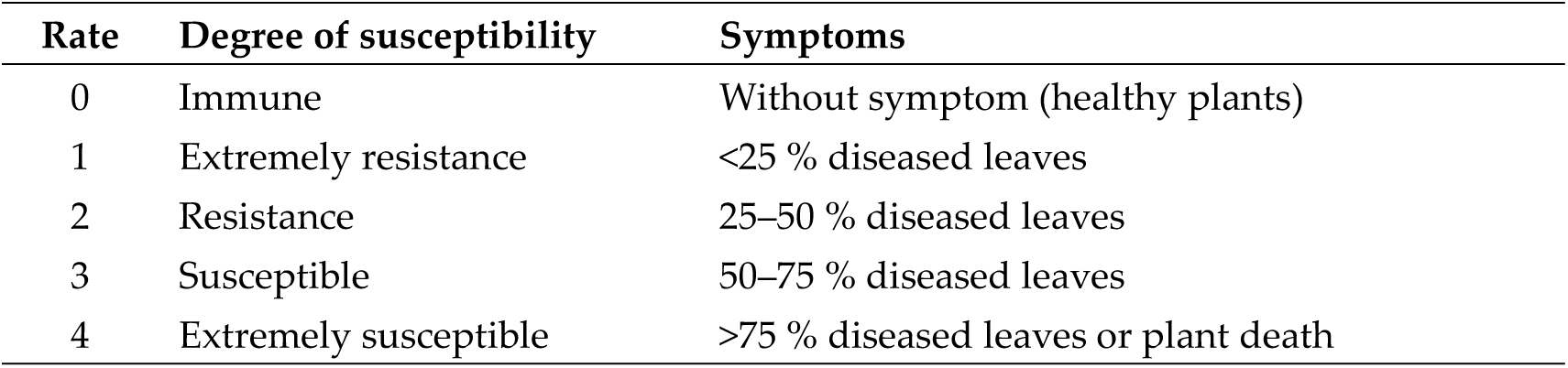
Scoring of symptoms of *Verticillium* wilt

| Rate | Degree of susceptibility | Symptoms |
| --- | --- | --- |
| 0 | Immune | Without symptom (healthy plants) |
| 1 | Extremely resistance | <25 % diseased leaves |
| 2 | Resistance | 25–50 % diseased leaves |
| 3 | Susceptible | 50–75 % diseased leaves |
| 4 | Extremely susceptible | >75 % diseased leaves or plant death |

The disease Index (DI) was estimated following the formulae below [7, 61].

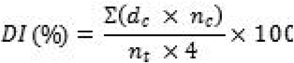

Where, *d_c_* is disease rate between 0 and 4;

*n_c_* is number of plants with interrelated disease rate;

*n_t_* is total number of plants tested for each CSSL

### 4.4 Analysis of phenotypic trait

The software SPSS 20.0 was used for analyzing the observed phonotypic data and the Pearson’s rank correlation coefficient was used for evaluating the correlation among the disease index. The statistical package SAS version 9.1was employed for Analysis of variance (ANOVA) of disease index and Tukey’s test was used to compare treatment means. The broad-sense heritability (H^2^) was calculated following the formulae described by [63].

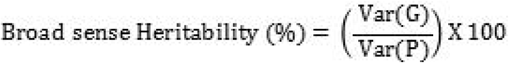

Where, Var(G) = Genotypic variance Var(P) = Phenotypic variance

### 4.5 Genetic analysis

Genomic DNA of CSSLs from BC_5_F_3:5_ population and its parents was extracted by following a modified procedure of CTAB method [64] by using young leaves which were sampled from each line and kept at −80°C. The working concentration of DNA was adjusted at 30ng/μL; quantified on NanoDrop2000 spectrophotometer (NanoDrop Technologies, Wilmington, DE USA). Further the integrity of DNA was patterned on agarose gel (1%) using Lambda DNA/HindIII Markers[65] as ladder. Scoring pattern followed for SSRs fragments include ‘–‘ for missing, ‘1’ for presence and ‘0’ for absence of bands.

### 4.6 SSR markers and SSR molecular detection

Based on the genetic map [37], in total, 597 pairs markers were screened out by using 2292 pairs of markers to be used to screen 300 CSSLs DNA. The sequences of these SSR primers were downloaded from the CMD database (http://www.cottongen.org/). First of all, we diluted these primer pairs. For dilution, we centrifuge primer pairs at 12000rpm at 4^0^C for 10 minutes to settle down the contents at the bottom. We diluted these primer pairs 100X and shake it vigorously for 2 minutes. Centrifuge it again and store at −20^0^C. The details of these SSR primers are mentioned in Table S2.

### 4.7 QTL mapping

QTL IciMapping V4.0 software developed by Wang et al. [66] was used to map QTLs of CSSLs. A LOD (likelihood of odds) of threshold 2.5 was used to declare significant additive QTLs. The percentage of phenotypic variance (PV%) explained individual QTL and additive effects at the LOD peaks were determined through stepwise regression (RSTEP-LRT). The graphical presentation of QTLs was done by using the MapChart2.2 software [67].

Positive additive effects showed that CCRI36 alleles decremented the phenotypic disease index values and enhanced resistance against VW. On the other hand, negative scores indicated that Hai1alleles decremented the phenotypic disease index values and incremented the values of VW resistance. The QTL nomenclature was designed as follows: the QTL designations begin with “q” come after the trait abbreviation, the chromosome name, and the number of QTL on that chromosome [68, 69]. Stable QTL was declared when it is found in at least two environments.

### 4.8 Meta-analysis of QTLs

Biomercator 4.2 [70] software was considered suitable for our data in order to perform Meta-analysis[32]. Already performed QTL meta-analysis has established a database[33]of QTLs including approximately 2,274 QTLs regarding 66 traits; accounting 201 QTLs regarding resistance for VW [13, 21, 43, 46–48, 50, 61, 71]. In our study, we kept the standard reference of Said et al [33]for information of mapped QTLs controlling VW resistance. Remaining previous studies, including 113 QTLs responsible for VW resistance have also been mentioned later[7, 19, 22, 45, 72, 73]. In aggregate 367 QTLs related to VW resistance have been utilized to build a platform for meta-analysis in which 53 QTLs were from our discovery in current study. On manual basis, new QTL hotspots have been identified by considering a consistent QTL region as if four or more QTLs were occurring in an interval of 25cM. However, the same consistent QTL region was possessing QTLs for only one trait then it was taken as a ‘QTL Hotspot’ [7].

Meta-analysis was performed by taking two files as input i.e. QTL file and map file. Map file was based on the information regarding names of parents, cross type and markers position on chromosomes. The QTL file was loaded with QTL in given environment as row information and QTL name, trait name, trait ontology, location, year, chromosome number, linkage group, LOD score, observed phenotypic variation (R^2^), most likely position of QTL, CI start position and CI end position. Initially, the two files were uploaded successfully and map connectivity was investigated for construction of consensus map. After that QTLs projection on consensus map was done, followed by meta-analysis regarding trait. Ultimately four model were obtained with different AIC (Akaike information criterion) value. The lowest AIC value holding model was considered suitable for the identification of mQTL positon or QTL hotspot. The criteria described by Said et al.[32]of occurrence of mQTLs in 20 cM interval was kept standard for the identification of hotspot.

## 5. Conclusions

In this study, 300 CSSLs developed from *Gossypium hirsutum* CCRI36 × *Gossypium barbadense* Hai1 were used to detect QTL for VW resistance in various environments (Anyang and Xinjiang) and different developmental stages (July and August). The nature of population (CSSL), population size and the presence of control (Jimian11) in our study showed us to lower the experimental error and to check the accurateness of data.

In total, 53 QTLs for VW resistance were identified in CSSLs populations, of which 29 ones were found as stable QTLs. Ten QTLs were similar to previously reported QTLs, while 43 ones were novel QTLs. Based on meta-analysis, 32 QTLs hotspot regions were detected, including 15 novel ones. These consistent QTLs and hotspot regions form critical steps, which will contribute to molecular breeders in developing and improving the VW resistance in upland cotton. The outcomes of this study also provide most important message for further studies of the molecular basis of VW resistance in cotton.

## Supplementary Materials

Supplementary materials can be found at www.mdpi.com/link.

## Ethics approval and consent to participate

Not applicable

## Consent for publication

Not applicable

## Author Contributions

Y.Y.L., T.T.C., Y.Z.S conceived and designed the experiments; M.H.R., L.P.T., K.K.P., Q.G., A.Y.L., J.W.G., Q.W.L., L.D., R.O.M., M.S.I., M.J. and W.K.G. performed the experiments; M.H.R. and L.P.T. analyzed the data; M.H.R. contributed reagents/materials/analysis tools: Y.Y.L., Y.Z.S., M.H.R., and L.P.T. drafted the manuscript.

## Abbreviations

English Abbr.: English Full Name
VW: *Verticillium* Wilt
DI: Disease Index
SSR: Simple Sequence Repeats
CSSL: Chromosome Segment Substitution Lines
AIC: Akaike Information Criterion
CI: Confidence Interval
Chr: Chromosome
cM: Centi-Morgan
CMD: Cotton Marker Database
H^2^_B_: Broad sense Heritability
LOD: Logarithm of Odds
MAS: Marker Assisted Selection
QTL: Quantitative Trait Loci
PV: Phenotypic Variation
CTAB: Cetyl-Trimethyl Ammonium Bromide
Mya: Million Years Ago

---

**Figure S2.**
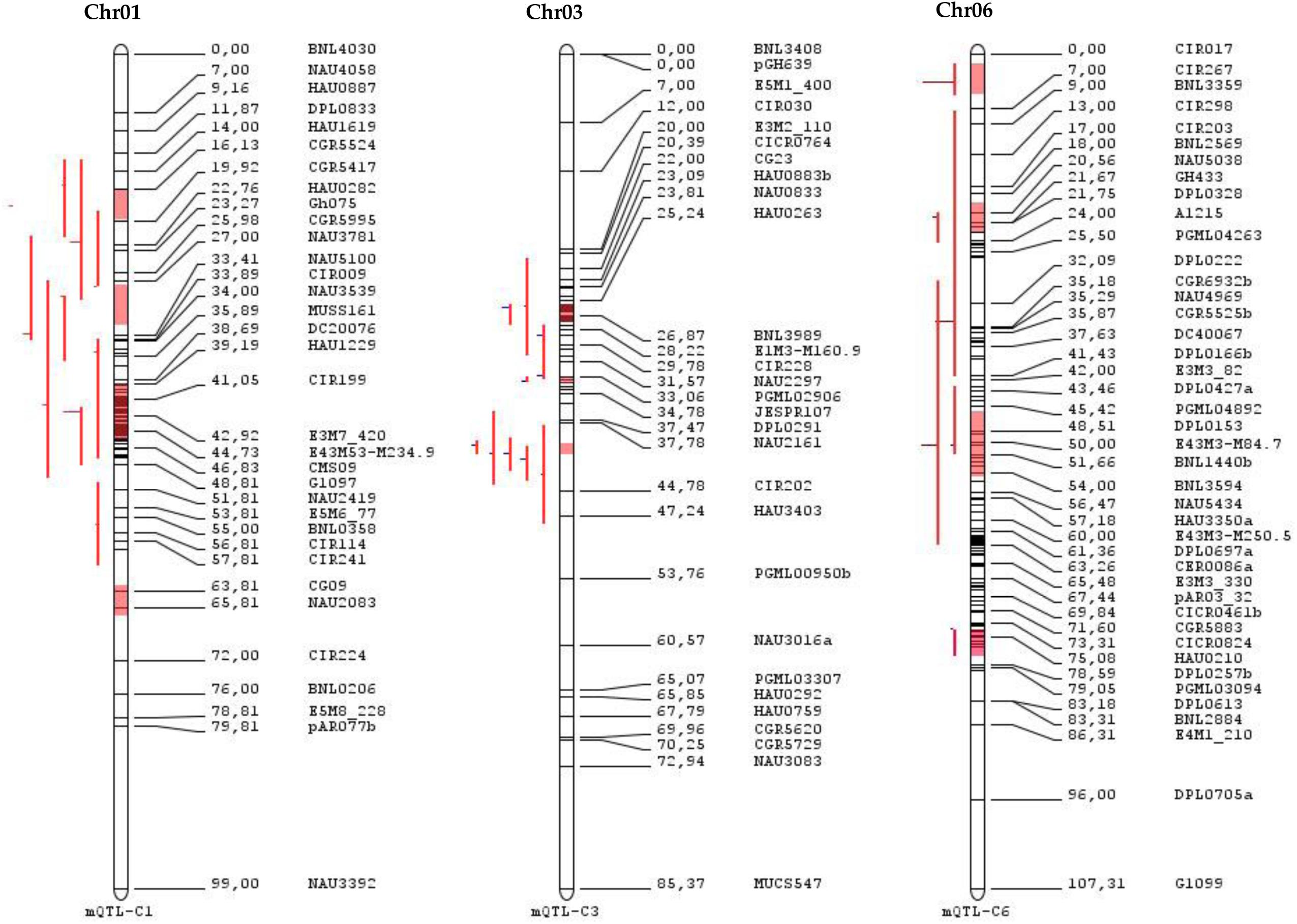
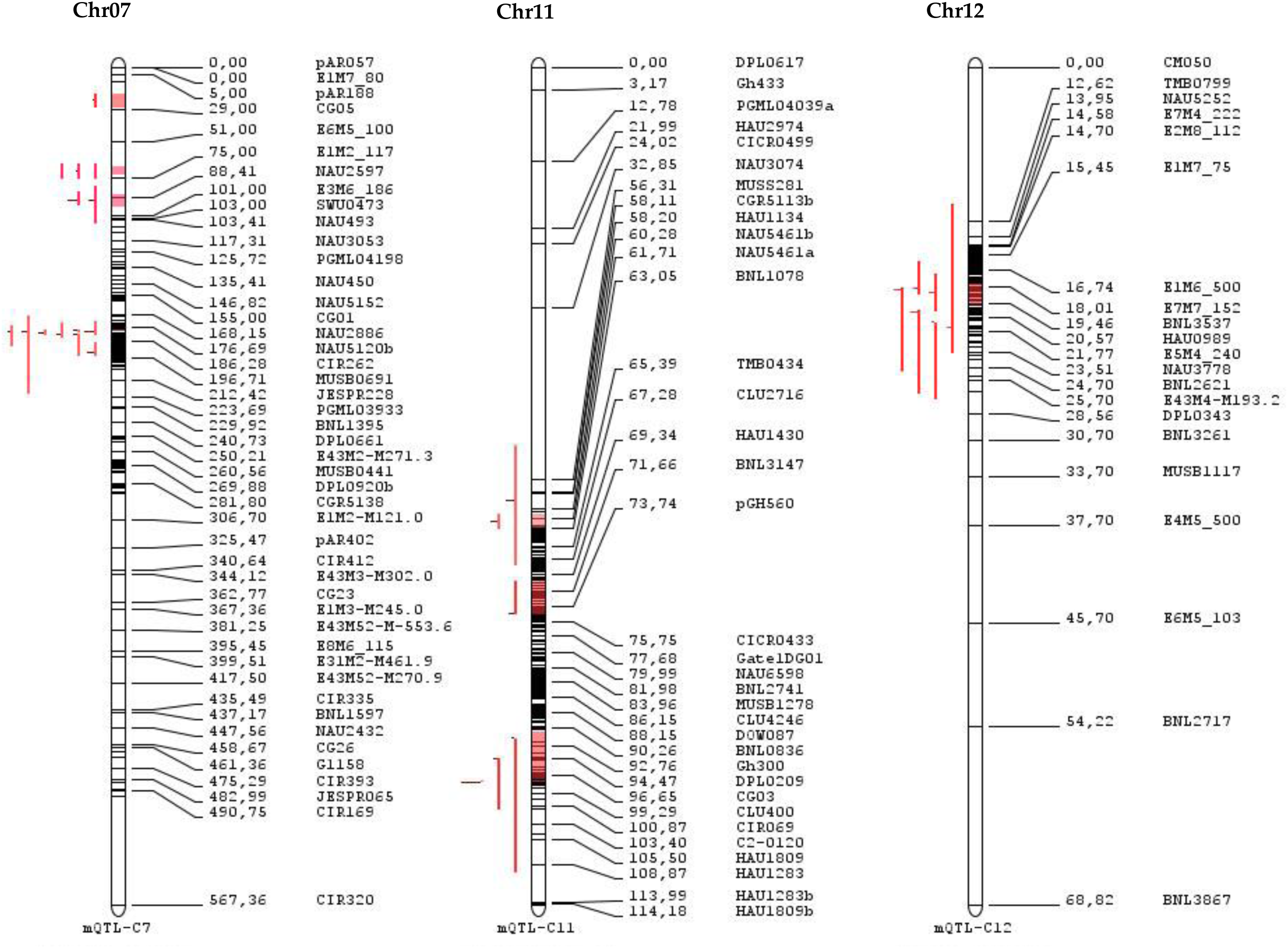
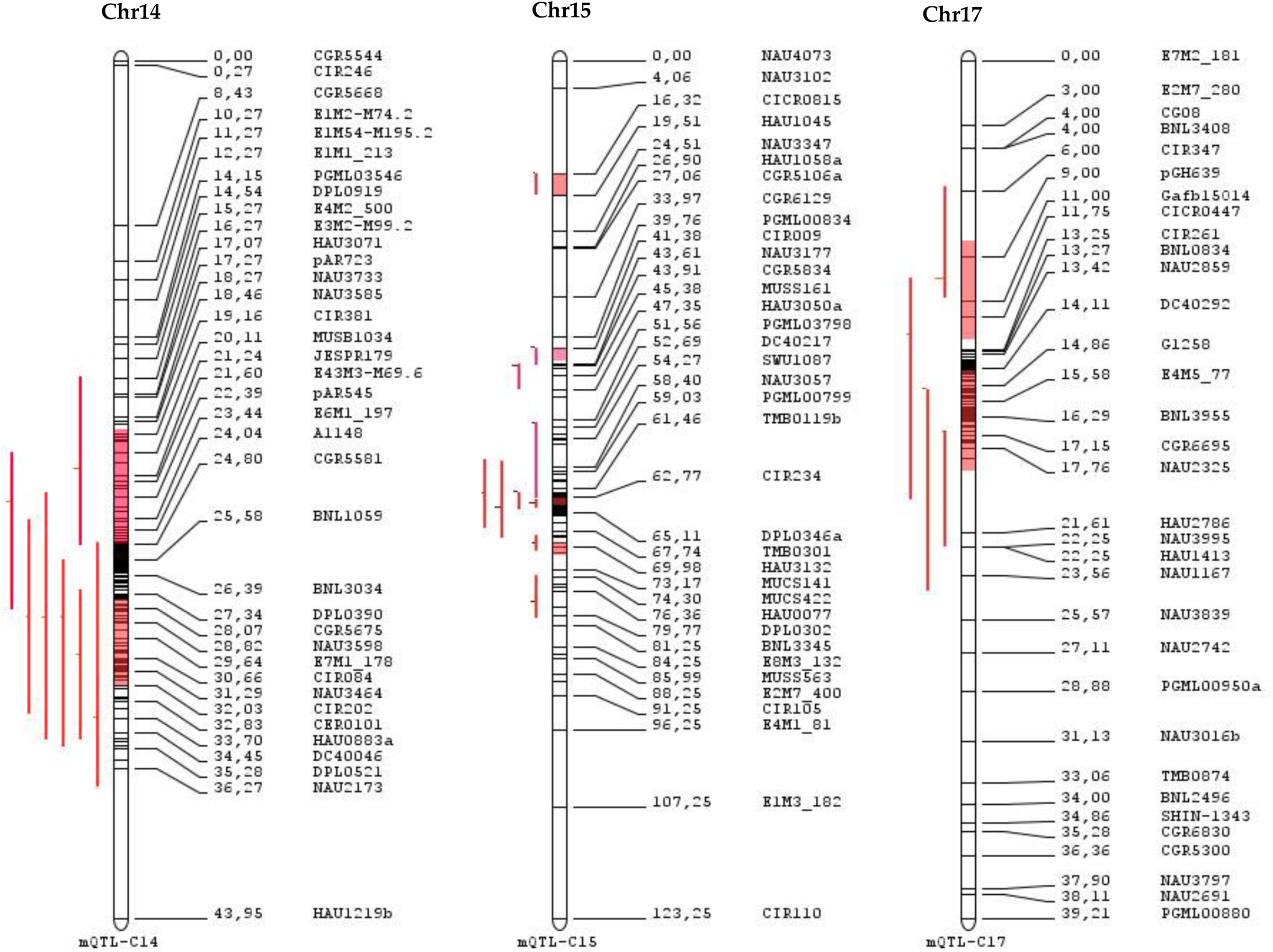
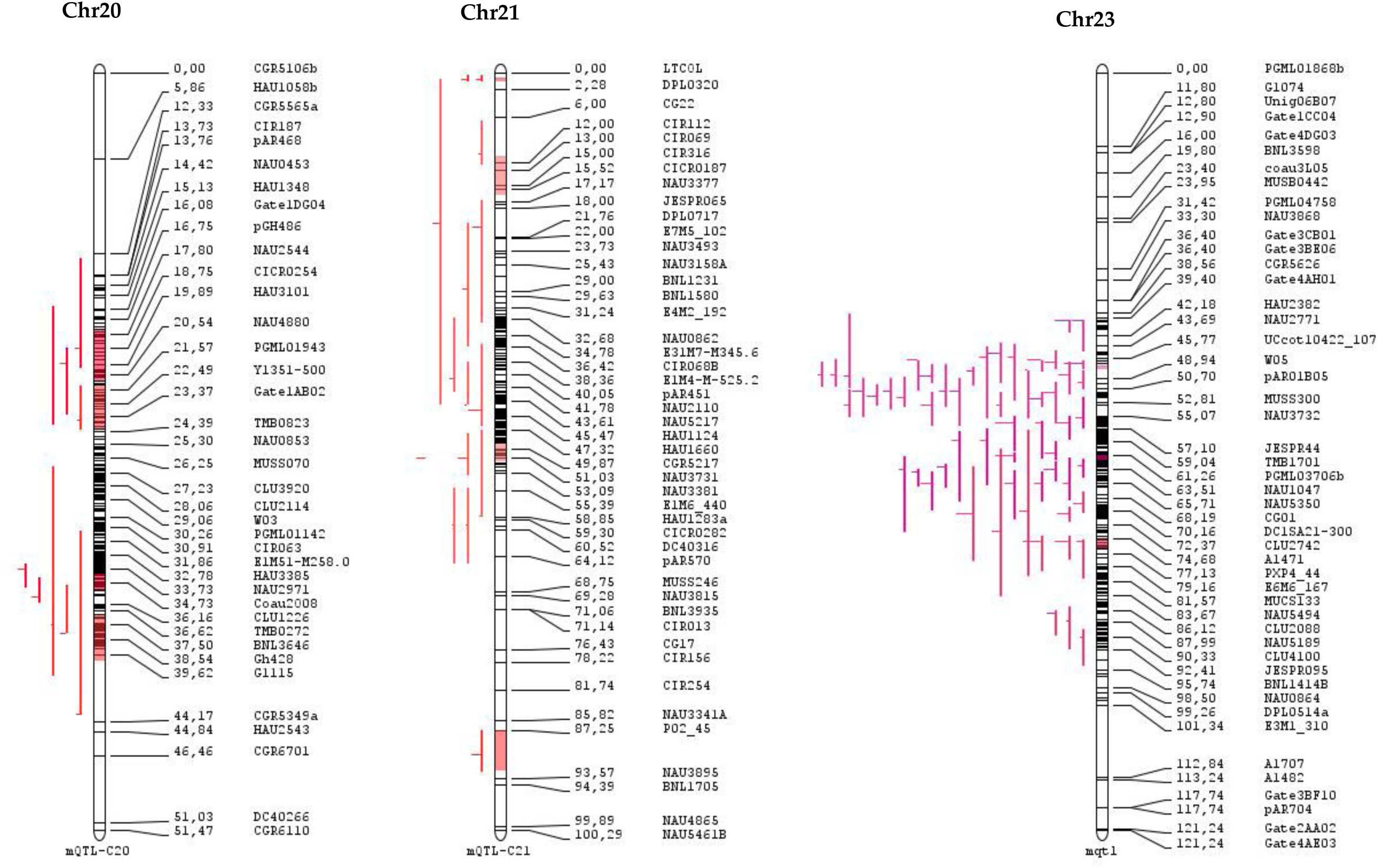
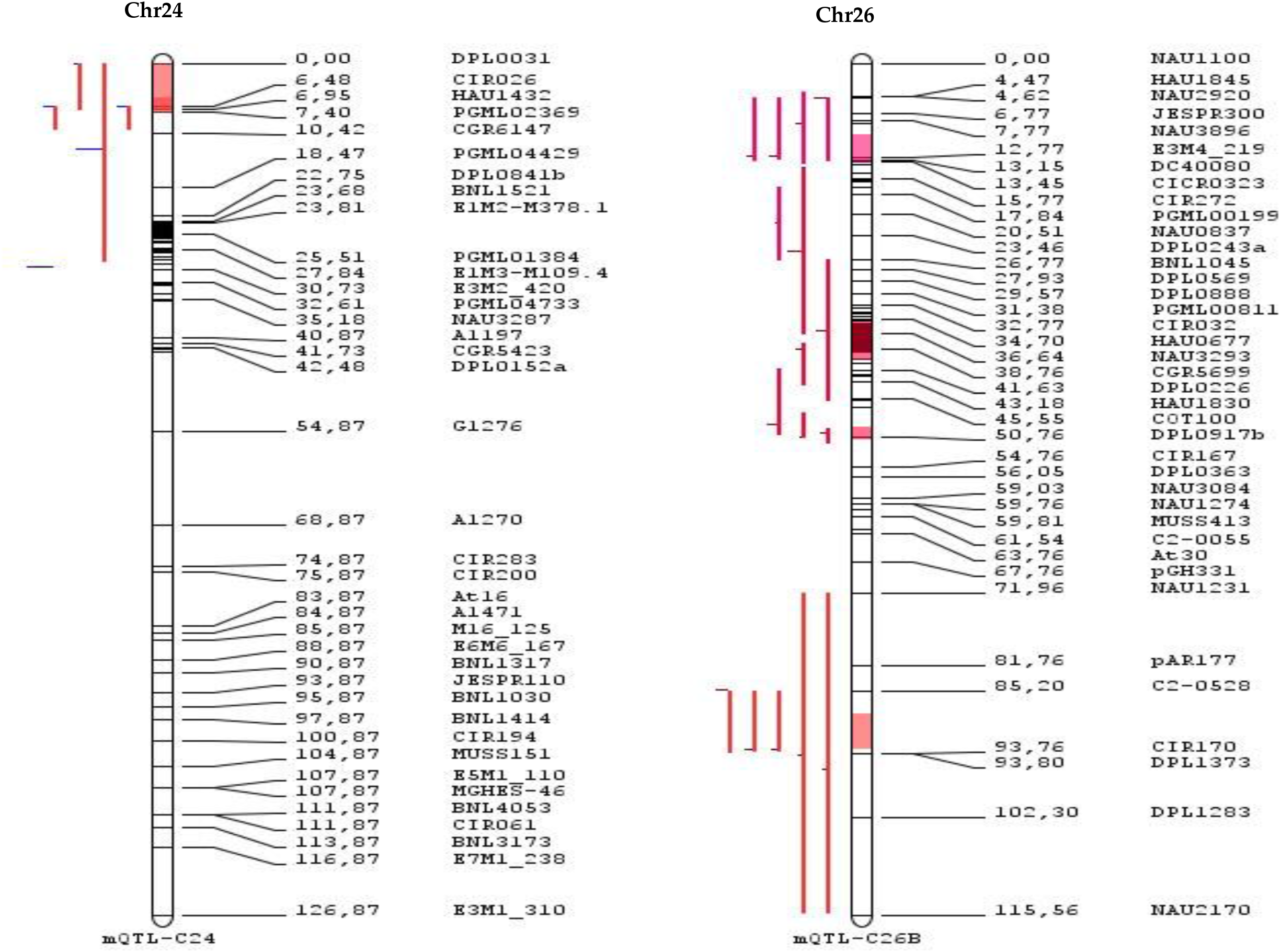
QTLs and QTL hotspot for *Verticillium* wilt resistance on the consensus map by a meta-analysis

**Table S1.**
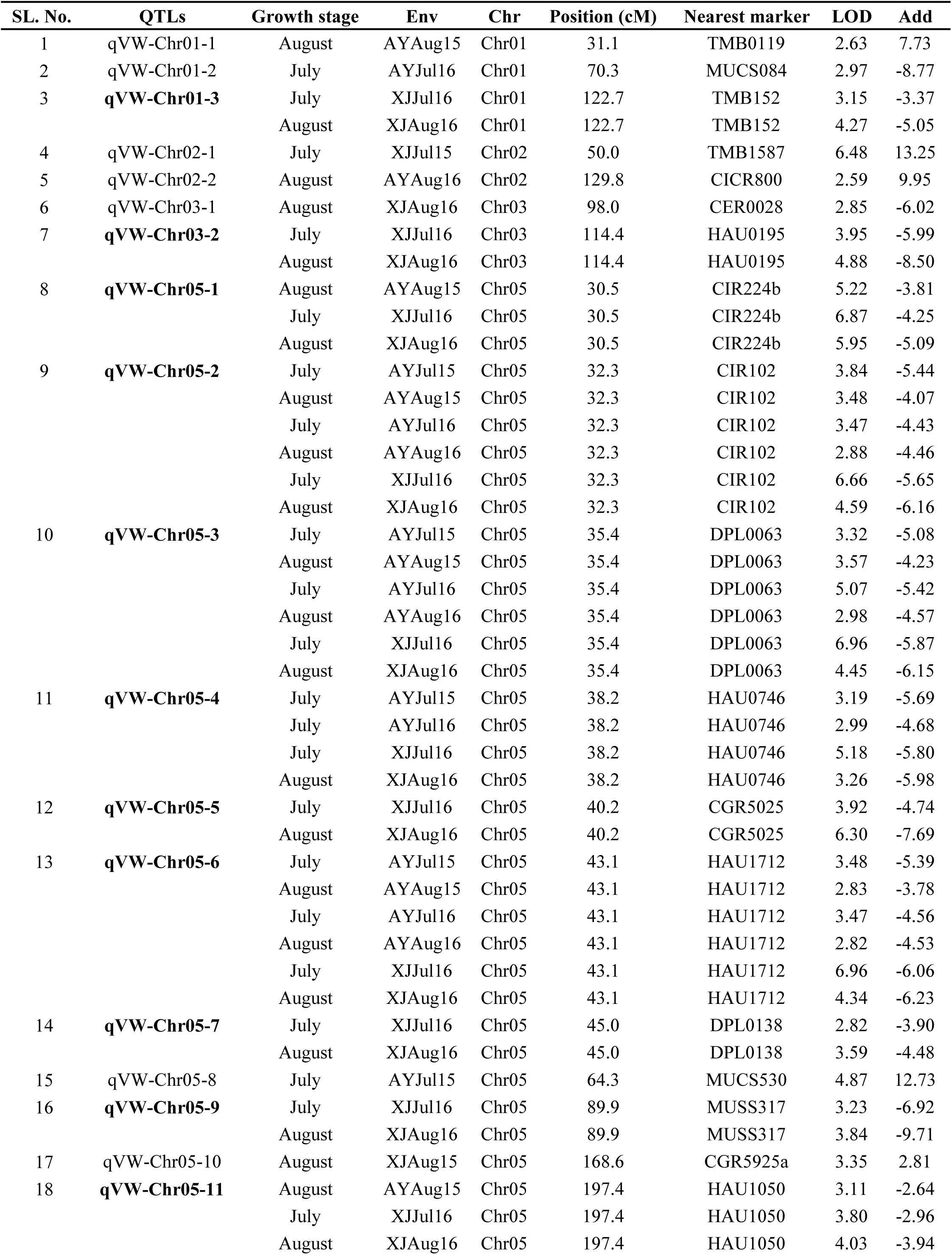
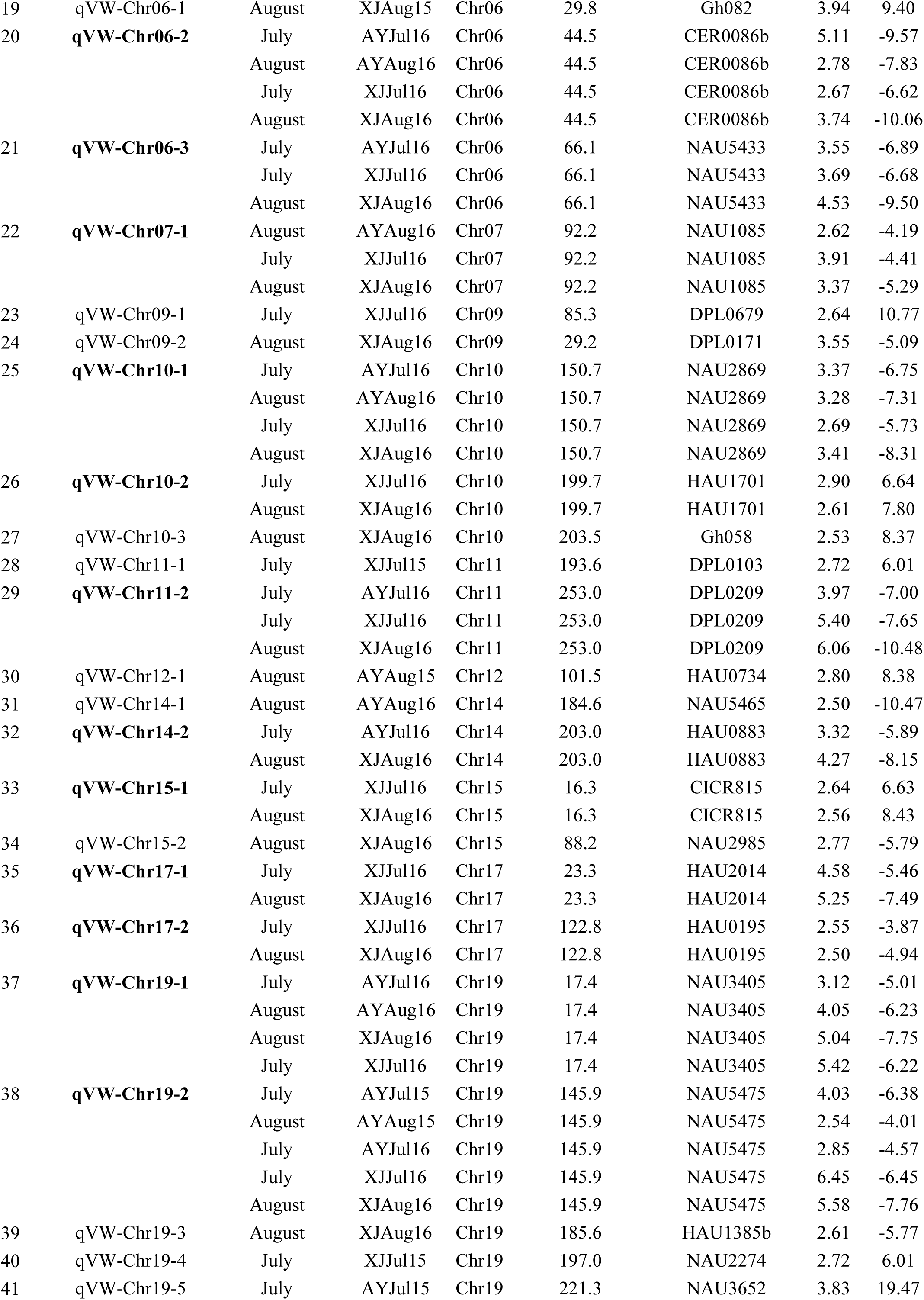
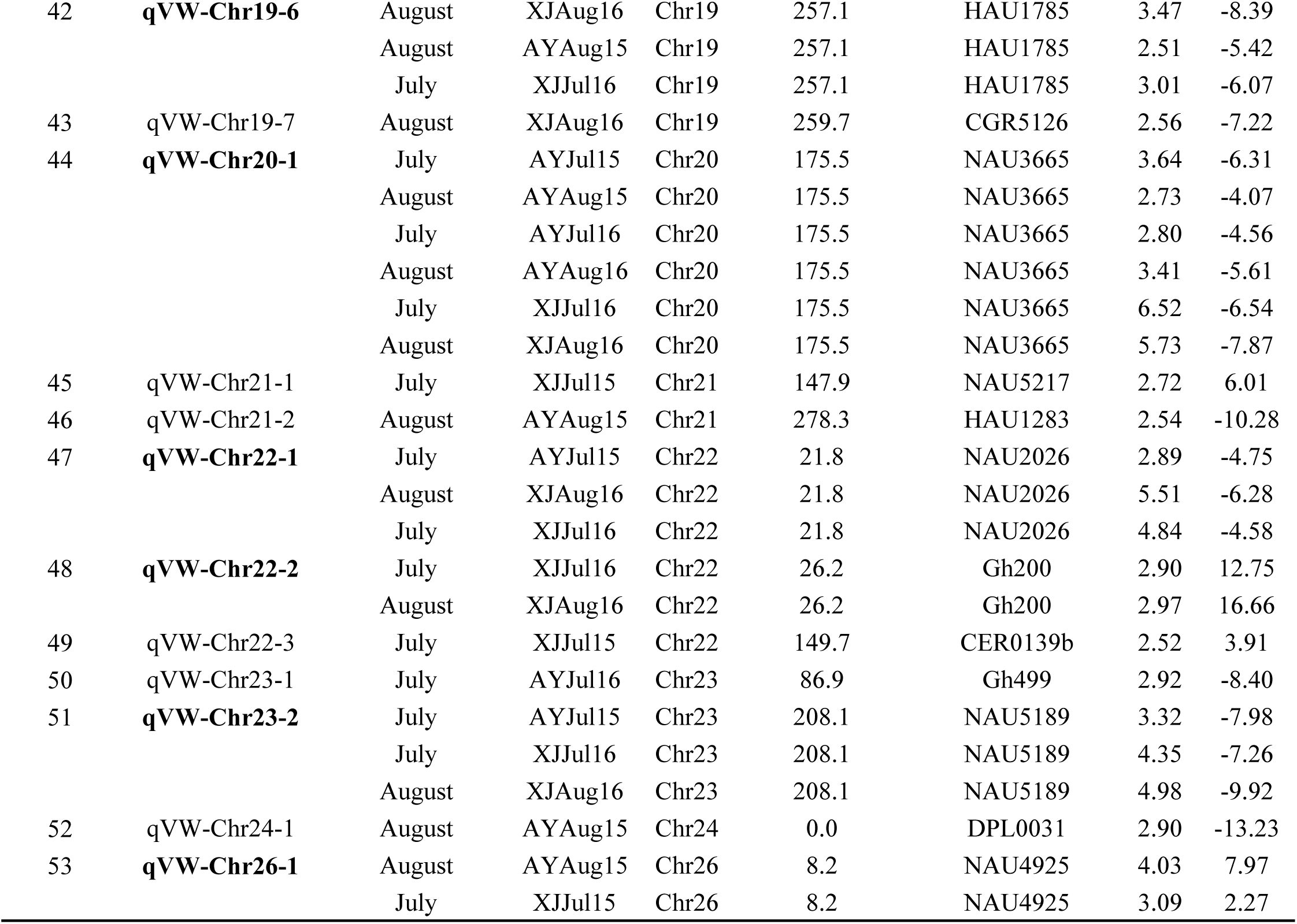
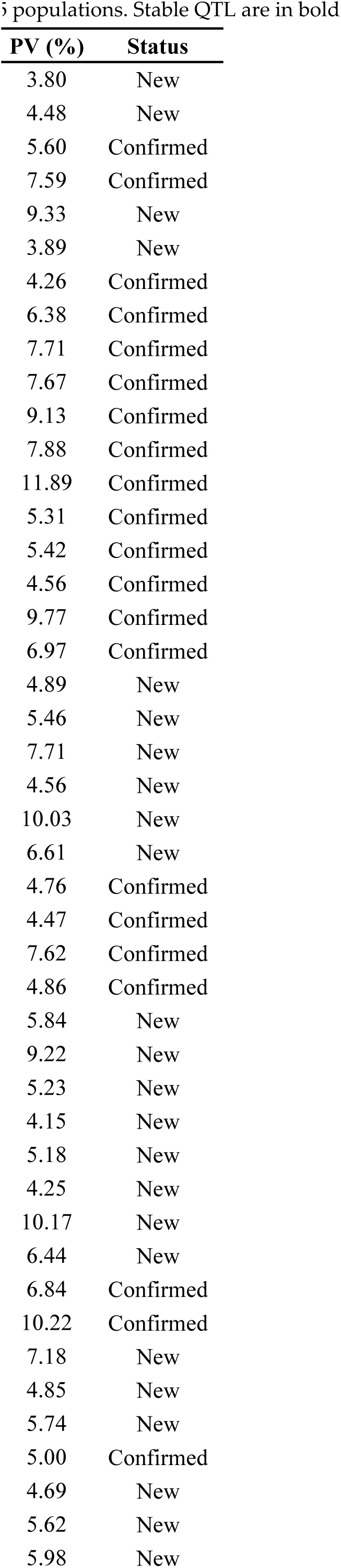
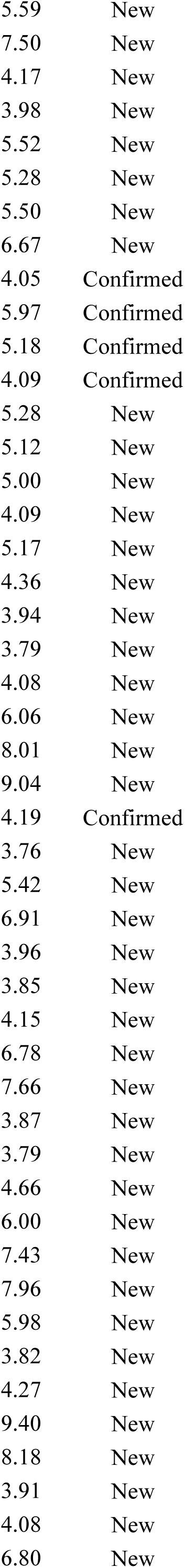
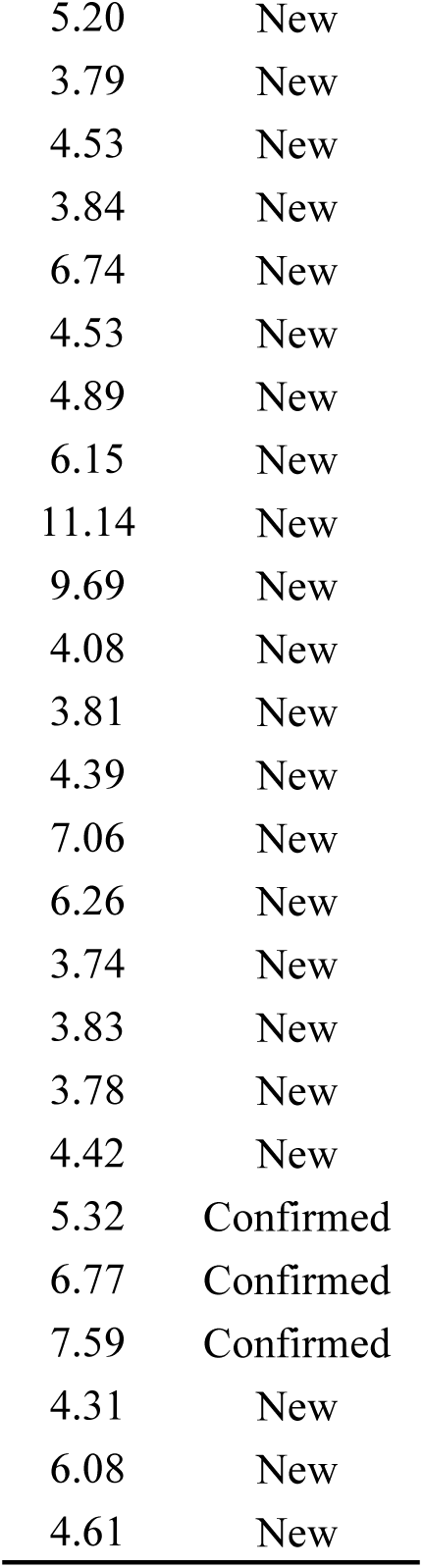
Details QTLs for Verticillium wilt resistance detected during different stages of growth and environments in BC5F3:5

**Table 2.**
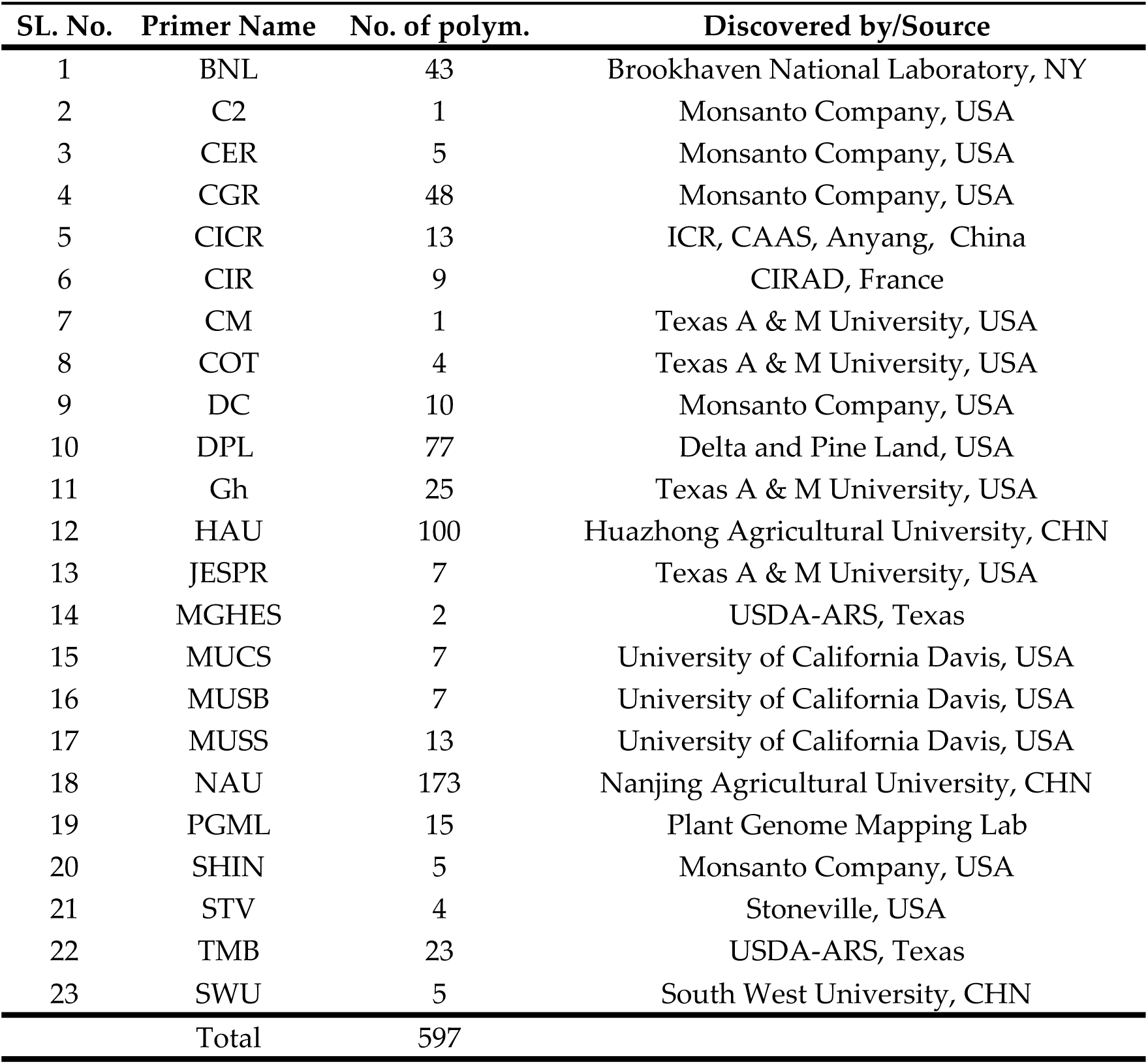
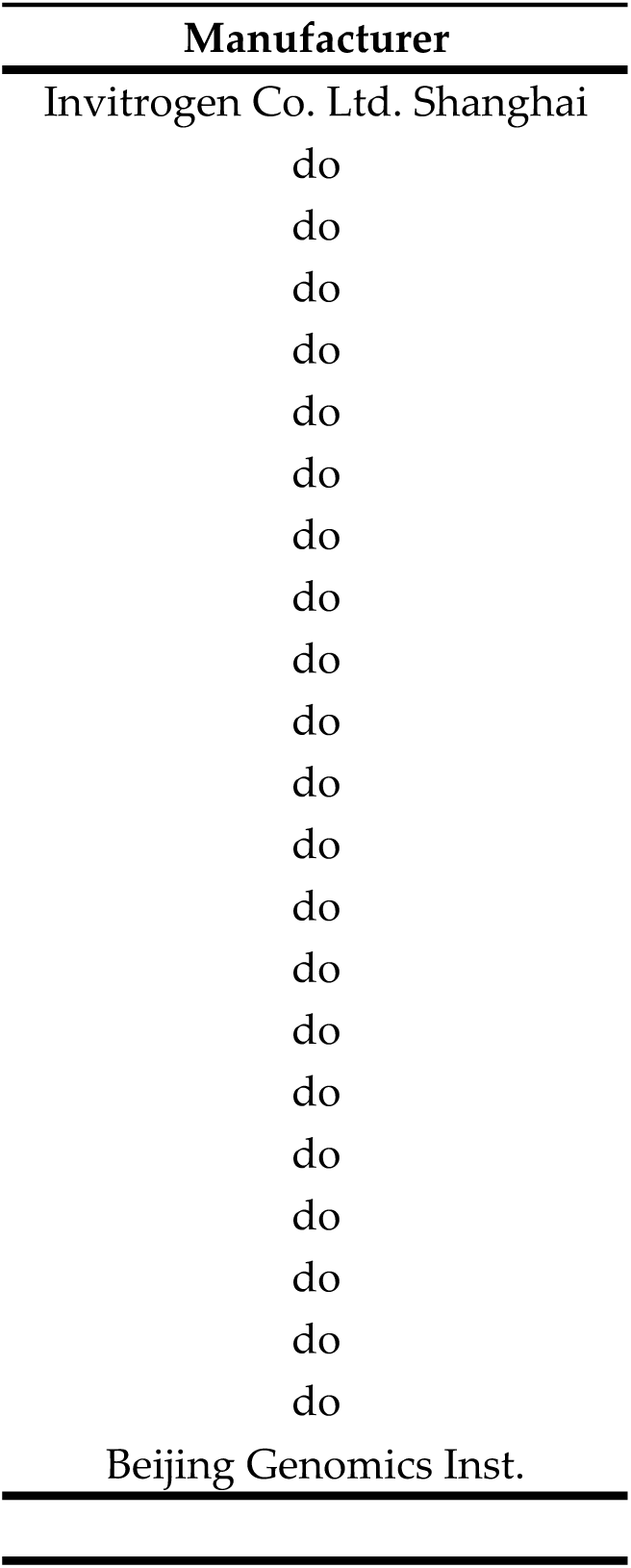
Details of primers used in this study

